# Identification of novel human 15-lipoxygenase-2 (h15-LOX-2) inhibitors using a virtual screening approach

**DOI:** 10.1101/2024.07.21.604444

**Authors:** Lucas Gasparello Viviani, Thais Satie Iijima, Erika Piccirillo, Leandro de Rezende, Thiago Geronimo Pires Alegria, Luis Eduardo Soares Netto, Antonia Tavares do Amaral, Sayuri Miyamoto

## Abstract

The human 15-lipoxygenase-2 (h15-LOX-2) is a non-heme iron-containing enzyme that catalyzes the regio- and stereospecific oxygenation of polyunsaturated fatty acids, mainly arachidonic acid, and is implicated in the biosynthesis of pro- and anti-inflammatory lipid mediators. The biological roles of h15-LOX-2 have not been completely unveiled, but it has been suggested that high expression levels of h15-LOX-2 are related to the pathogenesis of atherosclerosis and of some types of cancer. Inhibitors of h15-LOX-2 might be helpful for a deeper understanding of its roles in physiological and pathophysiological processes, in addition to representing potential drug candidates for treating human diseases. Nevertheless, only a few h15-LOX-2 inhibitors have been reported in the literature to date. Here, aiming to search for novel h15-LOX-2 inhibitors, we used a virtual screening (VS) approach, consisting of four consecutive filters (shape-based matching, 2D structural “dissimilarity”, docking, and careful visual inspection), which were applied to a “curated” version of the ZINC database, pre-filtered for potential drug-like compounds. Six novel h15- LOX-2 inhibitors, with inhibitory potencies in the micromolar range, were identified. K_i_ values were determined for two inhibitors, compounds **10** [K_i_ = (16.4 ± 8.1) μM] and **13** [K_i_ = (15.1 ± 7.6) μM], which showed a mixed-type mechanism of inhibition. According to docking predictions, the identified inhibitors occupy the more solvent-exposed arm of the U-shaped h15-LOX-2 active site’s cavity, possibly blocking the access of the substrate to the active site. The identified inhibitors are structurally different from the few h15-LOX-2 inhibitors reported in the literature, in addition to fulfilling drug-like criteria. Overall, our results provide a valuable contribution to the search for novel inhibitors of h15-LOX-2, a so-far underexploited target enzyme.

## INTRODUCTION

Lipoxygenases (LOXs) are non-heme iron enzymes that catalyze the regio- and stereospecific dioxygenation of polyunsaturated fatty acids containing at least two isolated cis double-bounds, such as arachidonic acid (C20:Δ4, n – 6) and linoleic acid (C18:Δ2, n – 6), leading to the formation of a chiral peroxide product.^1–4^ Humans have six LOXs (5-lipoxygenase, 12-lipoxygenase, 12R-lipoxygenase, 15-lipoxygenase-1, 15-lipoxygenase-2, and epidermal lipoxygenase-3), which are expressed in different tissues and named according to their product specificities.^1,5,6^ The primary products of the LOXs pathways are subsequently converted to several pro- or anti-inflammatory bioactive lipid mediators, including leukotrienes, lipoxins, hepoxilins, eoxins, resolvins, protectins, and others.^1,5^ In addition to playing important roles in the regulation of inflammation, LOXs are associated with biological processes such as cell proliferation and differentiation, modification of lipid-protein complexes, and regulation of the intracellular redox state.^1^ Additionally, LOXs have been implicated in the pathogenesis of many human diseases, including cancer, diabetes, and cardiovascular, pulmonary, and neurodegenerative diseases.^7^

The human 15-lipoxygenase-2 (h15-LOX-2), which is encoded by *alox15b* gene, converts arachidonic acid to 15(*S*)-hydroperoxyeicosatetraenoic acid (15(*S*)-HpETE) by attack at C13.^6^ h15-LOX-2 is expressed mainly in macrophages, skin, cornea, lungs, hair roots, and prostate.^8^ In epithelial cells, h15-LOX-2 has shown to regulate cell senescence.^9^ Recently, a role for h15-LOX-2 in macrophage cholesterol homeostasis has been described in the literature.^10^ Although h15-LOX-2 specific physiological roles have not been completely unveiled, it has been shown that h15-LOX-2 expression levels are higher in human carotid atherosclerotic lesions, in comparison to healthy arteries.^11–13^ Additionally, silencing *alox15b* gene in human macrophages has led to a decrease in cellular lipid accumulation and to a reduction in proinflammatory cytokine secretion.^14^ It has also been suggested that an increased expression of h15-LOX-2 induced by hypoxia may be associated to chemokine-mediated recruitment of T-cells in atherosclerosis and related diseases.^15^ Therefore, h15-LOX-2 seems to play an important role in the initiation and development of human atherosclerosis.^7,9^ In addition to its roles in atherosclerosis pathogenesis, several studies have suggested that h15-LOX-2 might be implicated in some types of cancer.^7^ h15-LOX-2 expression levels have shown to be altered in epithelial tumor cells^16–22^ and in tumor-associated macrophages from renal cell carcinoma.^23^ A possible role for h15-LOX-2 (and other LOXs) in ferroptosis, an iron-dependent form of cell death which is largely associated to lipid peroxidation, has also been proposed in the literature.^24–26^ However, the pharmacological importance of LOXs, including h15-LOX-2, as potential biological targets to ferroptosis-related diseases remains to be better studied.^24–26^

From a structural point of view, the available crystal structures of h15-LOX-2 (resolution values ranging from 2.44 to 2.63 Å)^6,8^ reveal that it has a cylindrical shape and displays a typical LOX fold, with an amino-terminal β-barrel domain (PLAT domain), which contains two Ca^2+^ binding sites, and a carboxy-terminal α-helical domain, which contains the substrate binding site and the catalytic iron (**Figure 1A**).^6,8^ The active site of h15-LOX-2 is in a U-shaped cavity formed primarily by hydrophobic amino acids (**Figure 1B**), which interact with the substrate non-polar fatty acid tail.^6^

**Figure 1.**
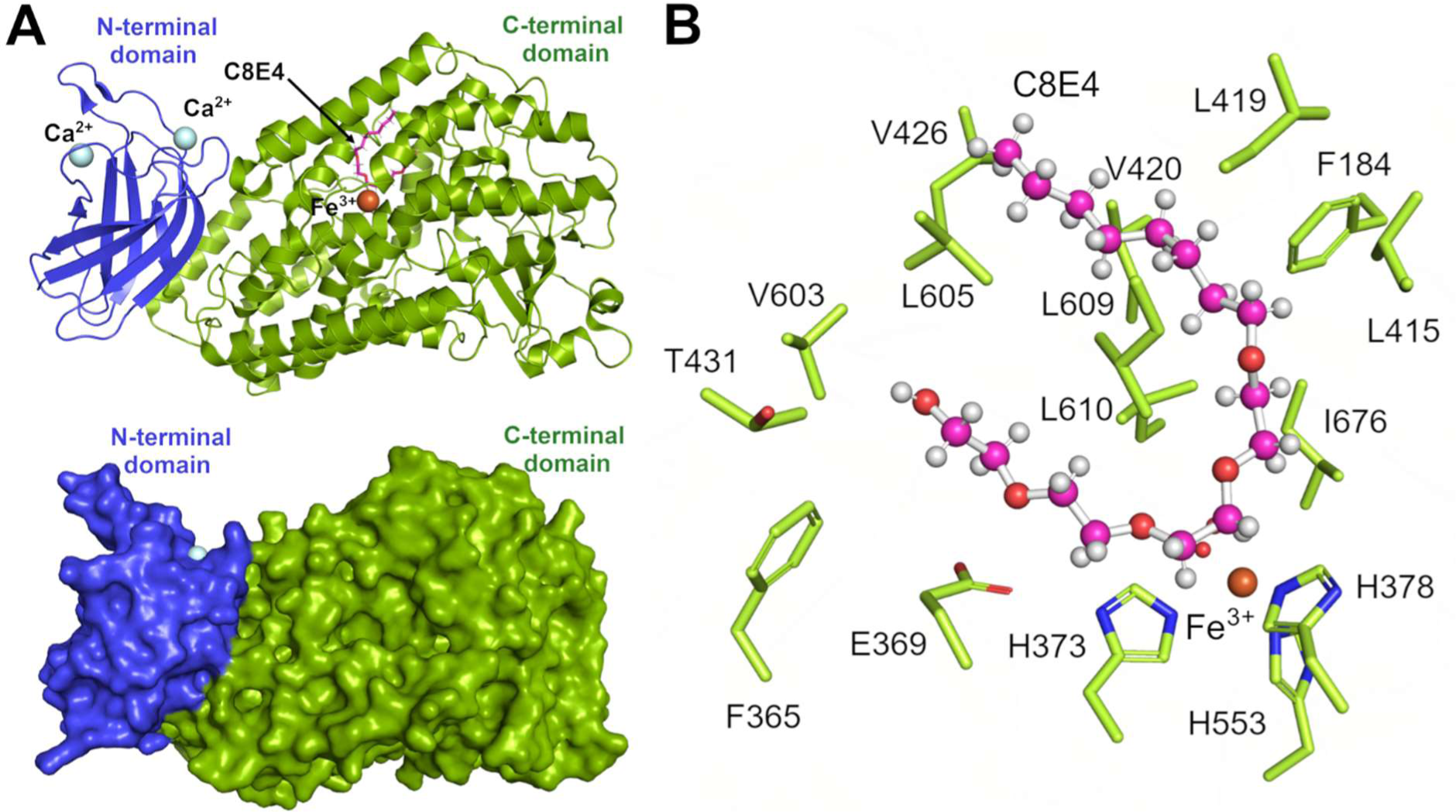
Representations of h15-LOX-2 overall structure and substrate binding site. **(A)** h15-LOX-2 structure in complex with a substrate mimic inhibitor (C8E4) (PDB code: 4NRE; resolution: 2.63 Å).^6^ Top: cartoon representation. Bottom: surface representation. The amino-terminal β-barrel domain (PLAT domain, the membrane-bound domain) and the α-helical carboxy-terminal domain (the catalytic domain) are shown in blue and in green, respectively. The catalytic iron (Fe^2+^/Fe^3+^) and two calcium (Ca^2+^) ions that play a role in membrane binding^6^ are shown as a brown sphere and as cyan spheres, respectively. The inhibitor C8E4 is represented as sticks (in pink). **(B)** h15-LOX-2 residues that form the active site. The residue side chains are shown as sticks (carbon and nitrogen atoms in green and in blue, respectively). The backbones and hydrogen atoms are omitted for clarity. The substrate mimic inhibitor (C8E4)^6^ is represented as ball-and-sticks (carbon, hydrogen, and oxygen atoms shown in pink, white and red, respectively). The catalytic iron is shown as a brown sphere. The figure was prepared using PyMOL.

Considering the complex (and so far poorly understood) biological functions of h15-LOX-2 and its potential as a therapeutic target for human diseases like atherosclerosis and cancer, discovering h15-LOX-2 inhibitors is of great interest for a deeper understanding of its roles in different physiological and pathophysiological contexts and/or for development of potential drug candidates.^6,8,27^ The elucidation of the h15-LOX-2 3D structure by X-ray crystallography^6^ has provided the basis for structure-based design of inhibitors for this target enzyme.^8^ Nevertheless, only a few h15-LOX-2 inhibitors have been reported in the literature so far.^6,8,27–31^ Among them, nordihydroguaiaretic acid (NDGA) and some flavonoid-based compounds have been described as moderately potent and non-selective inhibitors that act by reducing the active site’s catalytic iron.^28^ More recently, a series of low-micromolar h15-LOX-2 inhibitors that display some selectivity against other LOXs and cyclooxygenases (COXs) have also been reported.^8,27^ Still, some inhibitors of the h15-LOX-2/PEBP1 (phosphatidylethanolamine (PE)-binding protein 1) complex were found to inhibit the production of polyunsaturated fatty acid-phosphatidylethalonamine hydroperoxides, in addition to suppressing ferroptosis in cell culture models.^29^ However, the recently described inhibitors of h15-LOX-2 and of the h15-LOX-2/PEBP1 complex lack structural diversity since they are all derived from a similar scaffold, consisting of a central imidazole or oxadiazole ring substituted at two or three positions. Moreover, to our knowledge, the few known h15-LOX-2 inhibitors are still in preliminary stages of development as drug candidates and have not yet reached the clinics. Therefore, the need for discovering novel h15-LOX-2 inhibitors remains urgent.

Here, we used a virtual screening (VS) protocol, combining ligand-based^32^ and structure-based^32^ approaches to screen a “curated” version of the ZINC database^33,34^ (pre-filtered based on drug-like^35^ properties) for novel, structurally diverse and potentially drug-like h15-LOX-2 inhibitors. Six novel h15-LOX-2 inhibitors, with inhibitory potencies in the micromolar range, were identified through *in vitro* enzyme inhibition assays for VS experimental validation. The identified inhibitors are structurally diverse from the h15-LOX-2 inhibitors reported in the literature so far,^6,8,27–31^ show drug-like properties, and were predicted to interact with the more solvent-accessible arm of the U-shaped active site cavity.

## MATERIALS AND METHODS

### Protein structure preparation for the virtual screening procedures

The h15-LOX-2 structure, in complex with a substrate mimic inhibitor (C8E4), was taken from the Protein Data Bank (PDB code: 4NRE; resolution: 2.63 Å).^6^ and prepared for the docking procedures using SYBYL-X^36^ v.2.1. All hydrogen atoms were added and the protonation states of the ionizable amino acid side-chain groups were set at pH = 7.4. The side chain amide groups of Asn and Gln residues were oriented in the direction of maximal potential hydrogen bonding. All water molecules, as well as the co-crystallized ligand molecule (substrate mimic inhibitor), were removed from the original pdb file. The coordinates of two water molecules that coordinate iron were extracted and considered for docking procedures, as described below. The catalytic iron and two calcium ions were kept as part of the protein structure. An energy minimization of the protein structure in successive stages was performed with the standard TRIPOS force field, using the Powell’s method^37^ with simplex initial optimization, gradient termination of 0.05 kcal.mol^-1^.Å^-1^, and the maximum of iterations set to 500.

### Compounds database preparation

The structures of compounds from the ZINC-12^33,34^ database (∼23 x 10^6^ compounds) were downloaded in sdf format. Compounds from this database were submitted to a preliminary “curation” step, using the following criteria: (*i*) compounds should have good predicted water solubilities based on their calculated “Solubility Forecast Index” (SFI)^38^ values (compounds with SFI values ≤ 5.0 were considered to have a good predicted water solubility); and (*ii*) compounds should match the physicochemical and structural properties criteria defined according to the “drug” filter^39^ implemented in FILTER^40^ v.2.1.1 (with some modifications), which include: not violating more than two of the Lipinki’s Rule of Five^41,42^ criteria; having at least 15 non-hydrogen atoms; having 2-20 “heteroatoms” (non-carbon, non-hydrogen atoms); having a maximum of four chiral centers; having a maximum of 20 rotatable bonds; not having “promiscuous” protein-reactive groups^43,44^ (*e.g*., quinones, aldehydes, Michael acceptors, epoxides, acid halides etc.); among others. Compounds that passed the filter criteria (∼8 x 10^6^) compose a database referred to hereafter as “ZINC-Curated”. Compounds from the ZINC-Curated database were prepared for the VS procedures as follows: (*i*) the most abundant tautomeric forms and protonated states at pH = 7.4 were obtained using MoKa^45^ v.2.6; and (*ii*) up to 30 conformations per compound were generated using OMEGA^46–48^ v.2.5.1, with default parameters.

### Shape-based model generation and screening procedures

Two known h15-LOX-2 inhibitors reported in the literature (referred to as MLS000536924 and MLS000545091)^27^ were used to generate a shape-based model (“query”), using ROCS^49,50^ v.3.4.3.0. The 3D structures of the two inhibitors were obtained in sdf format using OpenBabel^51^ v.2.3.1 and, subsequently, the most abundant tautomeric form and protonation states of each compound at pH = 7.4 were obtained using the *Tautomers* and *FixpKa* tools from QUACPAC^52^ v.2.0.0.3. Next, a low energy conformation of each inhibitor was obtained using OMEGA^46–48^ v.2.5.1.5. Finally, the two compounds were aligned, using LigandScout’s^53^ v.4.4.7 alignment algorithm. The aligned compounds were used to generate a shape-based model (“query”) in ROCS, and default data settings were used to run a shape-based screening in ROCS. As described in the literature,^50^ ROCS performs overlays of conformers of the candidate molecule to the generated “query” based on their shape-matching. For each compound in the database, the conformer that best matched the model was ranked according to its Shape Tanimoto^50^ score value, which is computed based on the shape overlap and ranges from 0 to 1.

### 2D Structural “dissimilarity”^54^ filter

To eliminate compounds that showed a high degree of structural similarity among each other, compounds were first clustered by 2D structural similarity, using molecular fingerprints, which are binary sets (“bits”) that encode the presence/absence of a certain substructure/fragment in a molecule.^54^ The molecular fingerprints were obtained using the Open Babel program.^51^ Based on the fingerprints generated, the value of Tanimoto’s coefficient^55^ (TC) was calculated for each pair of compounds. The “dissimilarity” indices (DI) were then calculated from the Tanimoto’s coefficient values, as follows:

DI*_AB_* = 1 − TC*_AB_*, in which:
DI*_AB_*: dissimilarity index for two molecules, *A* and *B*.
TC*_AB_*: Tanimoto’s coefficient index for two molecules, *A* and *B*.

The calculated DI values were transferred to a data matrix, which was used as an input file in the R program^56^ for grouping structurally similar compounds into clusters, using a hierarchical clustering algorithm with complete linkage and the *hclust* function. Next, a DI cutoff value of 0.15 was applied, which means that compounds showing more than 85% of 2D similarity to each other were grouped into the same cluster. Only one representative compound of each cluster was maintained for the next VS steps.

### Docking and visual inspection procedures

Docking was performed using GOLD^57^ v.5.2. The h15-LOX-2 structure (PDB code: 4NRE; resolution: 2.63 Å) was prepared as described above and used for the docking calculations. The active site’s cavity, which was taken as the binding site region for docking, was defined by the residues Phe-184, Glu-369, His-373, His-378, Leu-415, Leu-420, Val-426, Leu-605, Ala-606, Leu-609, Leu-610, Ile-676, and by the catalytic iron. Two water molecules, which coordinate the protein’s catalytic iron, were allowed to switch on and off (*i.e*., protein-bound or displaced by the ligand) according to GOLD’s estimations of free-energy changes associated with moving a water molecule from the bulk solvent to the protein’s binding site. Ten docking runs were performed for each ligand, using default settings for genetic algorithm parameters, and ChemPLP^58^ and Goldscore^57^ as scoring functions. Docking procedure was validated by “re-docking” the substrate mimic inhibitor C8E4 (co-crystallized with h15-LOX-2)^6^ into the active site’s cavity, using the same settings as described above, except for not considering the two coordinating water molecules. A careful visual inspection analysis of the best-scored pose of each docked compound was performed, using LigandScout^53^ v.4.4.7 and PyMOL^59^ v.2.1.0, to check the fit into h15-LOX-2 active site’s cavity. In this analysis, the overall quality of the binding was evaluated based on recognition of the main protein-ligand interactions and on the shape complementarity between the compound and the protein’s binding site. Docking poses that showed unrealistic, sterically hindered conformations were excluded.

### h15-LOX-2 expression and purification

*E. coli* BL21 DE3 pLysS cells were transformed with pETDuet-1 containing the gene coding His-tagged h15-LOX-2 (50-60 ng), by electroporation. After transformation, cells grew up overnight as cultures in LB broth containing ampicillin (100 μg.mL^-1^) and chloramphenicol (170 μg.mL^-1^), at 37 °C, for ∼20h, at 150-190 rpm, and were subsequently diluted with 500 mL of LB medium to a concentration enough to initiate cell growth with an OD_600 nm_ = 0.2. Cells were maintained at 37 °C under agitation, until the exponential growth phase was reached (OD_600 nm_ = 0.6-1.0). Gene expression was then induced with isopropyl 1-thio-β-D-galactopyranoside (IPTG, 0.5 mM), in LB broth, for 12 hours, at 20 °C, under agitation (170 rpm). Afterward, cells were centrifuged (5000 rpm, 4 °C, 15 minutes) and resuspended in Tris-HCl buffer (50 mM, pH = 8,0) containing NaCl (500 mM), lysozyme (50 µg.mL^-1^), and protease inhibitor (1× *Sigma FAST Protease Inhibitor Tablet*). Cells were lysed in an ice bath by sonication for 5 minutes (40% amplitude, alternating between 20 s of sonication and 60 s of rest). Next, streptomycin (10% w/w) was added, and the lysate was maintained in ice by 15 minutes, under agitation. The lysate was then centrifuged (14,000 rpm, 4 °C, 40 minutes), and soluble proteins were filtered using a Durapore 0.45 μm PVDF filter (*Merck*, Germany).

h15-LOX-2 was purified by affinity chromatography (Ni^2+^-NTA Superflow, 1 mL), using the ÄKTA Protein Purification system (*GE Health Care Life Sciences*). The column was first washed (“equilibrated”) with 10 mL of purified water and with 10 mL of “Buffer A” (Tris-HCl (50 mM, pH = 8) and NaCl (500 mM)), with a flow rate of 1 mL.min^-1^. Purification was performed in four steps, with a flow rate of 1 mL.min^-1^, and the protein was eluted with a 5-100% gradient of “Buffer B” (Tris-HCl (50 mM, pH = 8), NaCl (500 mM) and imidazole (500 mM)). Fractions were analyzed by SDS-PAGE and those containing h15-LOX-2 were pooled and concentrated with an Amicon Ultra Centrifugal Filter (30 kDa cutoff) (*Merck*, Germany). Finally, a HiTrap™ 5 mL Desalting (*Cytiva*, USA) column was used for removing imidazole. The concentration of purified h15-LOX-2 was estimated by absorbance measurements at λ = 280 nm (ε = 115,445 M^-1^.cm^-1^),^60^ at room temperature, in a quartz cuvette.

### Enzymatic assays

All enzymatic assays were carried out in quartz cuvettes (path length of 1 cm), in a total volume of 300 μL, containing Tris-HCl buffer (25 mM, pH = 7.4), NaCl (250 mM), h15-LOX-2 (120-420 nM), Triton X-100 (0.014% v/v), arachidonic acid (25 μM), and the tested compound (1.0 μM to up to 199.1 μM, depending on the compound solubility in the assay buffer). The system was pre-incubated at room temperature for 2 minutes with all reagents, except for the substrate (arachidonic acid). Next, the enzyme reaction was started by adding arachidonic acid. The enzyme reaction was monitored spectrofotometrically, following the formation of the product ((5Z,8Z,11Z,13E)-15(*S*)-hydroperoxyeicosa-5,8,11,13-tetraenoic acid, 15(*S*)-HpETE; ε = 25,000 M^-1^.cm^-1^) at λ = 234 nm, for 20-100 seconds, at room temperature. Stock solutions of the tested compounds were prepared in DMSO, and the concentration of DMSO in the assays was 1.0% (v/v). Each assay was performed at least in duplicates. Data analyzes were carried out using the program GraphPad Prism v.9. A four-parameter logistic nonlinear regression model was used to build the dose-response curves and to obtain the IC_50_ values. A mixed-type inhibition model was used to calculate K_i_ values, after inhibition type diagnosis based on Lineweaver-Burk plots.

## RESULTS AND DISCUSSION

An overview of the hierarchical VS protocol that was used in this work to search for novel h15-LOX-2 inhibitors is schematically represented in **Figure 2A**.

**Figure 2.**
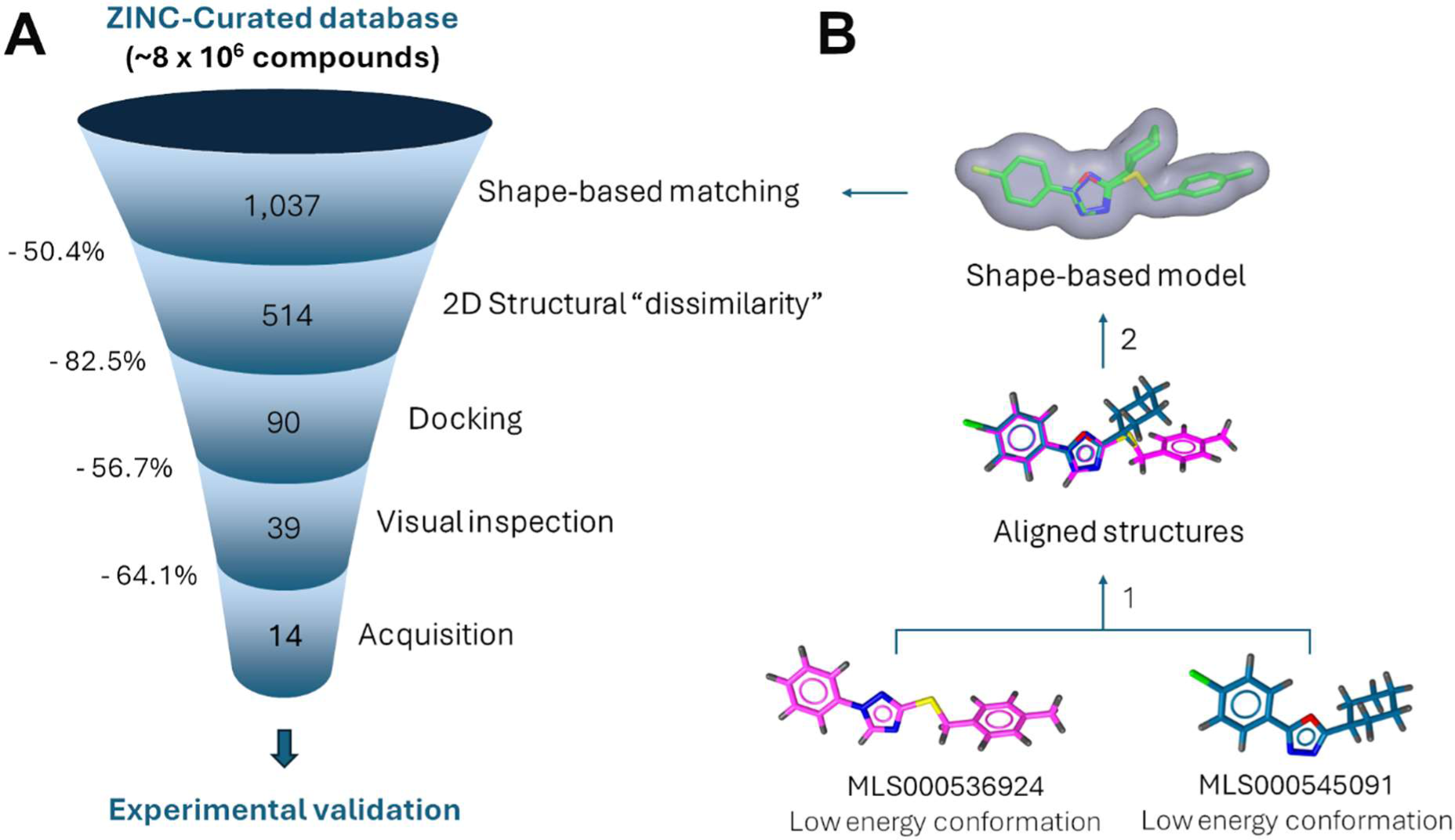
Schematic representation of the virtual screening (VS) protocol used in this study to select novel potential h15-LOX-2 inhibitors from the ZINC-Curated database. **(A)** Representation of the sequence of selection filters applied to the ZINC-Curated database in each VS step. **(B)** Schematic representation of the steps to generate the shape-based model that was used as a first filter in the VS protocol: 1. Alignment of low energy conformation structures of two known h15-LOX-2 inhibitors (MLS000536924 and MLS000545091),^27^ using LigandScout’s alignment algorithm; and 2. Shape-based model generation, using ROCS. The molecular shape surface is represented in gray. The figure was prepared using LigandScout and ROCS.

### Shape-based screening

As discussed in the literature, shape complementarity is one of the most important aspects of protein-ligand recognition,^50,61,62^ mainly for lipid substrates, which usually bind large, buried, and hydrophobic binding sites inside their target proteins.^63^ This is the case of arachidonic acid U-shaped binding site in h15-LOX-2, which is located in a highly hydrophobic, neutral, and deep cavity, as revealed by lipophilic potential, electrostatic potential, and depth cavity maps generated using SYBYL (**Figure 3**). Motivated by the recognized importance of shape complementarity for protein-ligand binding, as well as by the successful applications of shape-based approaches in the search for inhibitors of membrane-bound receptors and enzymes that bind lipids (see, *e.g*., references ^64–71^) and other types of substrates,^72^ we used a shape-based matching approach as a first filter in our VS protocol for selecting novel h15-LOX-2 inhibitors (**Figure 2A**). Two known and selective h15-LOX-2 inhibitors reported in the literature (referred to as MLS000536924 and MLS000545091),^27^ which probably bind the enzyme’s active site and show inhibition in the micromolar range, were used to build a shape-based model (“query”),^50^ using ROCS^50^ (**Figure 2B**). At the time our shape-based model was proposed, there were no crystal structures (neither other experimental data) that could give us a clue on the possible binding modes of MLS000536924 and MLS000545091 to h15-LOX-2. This was not a drawback, however, since the knowledge of the bioactive (experimentally determined) conformation(s) has not shown to significantly impact ROCS^50^ screening performance, as discussed in the literature.^50^ To circumvent the lack of information on the bioactive conformation(s) of MLS000536924 and MLS000545091, a low energy conformation was generated for each of these inhibitors, using OMEGA.^46–48^ The structures of MLS000536924 and MLS000545091 were subsequently aligned to each other and used to build a shape-based model (**Figure 2B**).

**Figure 3.**
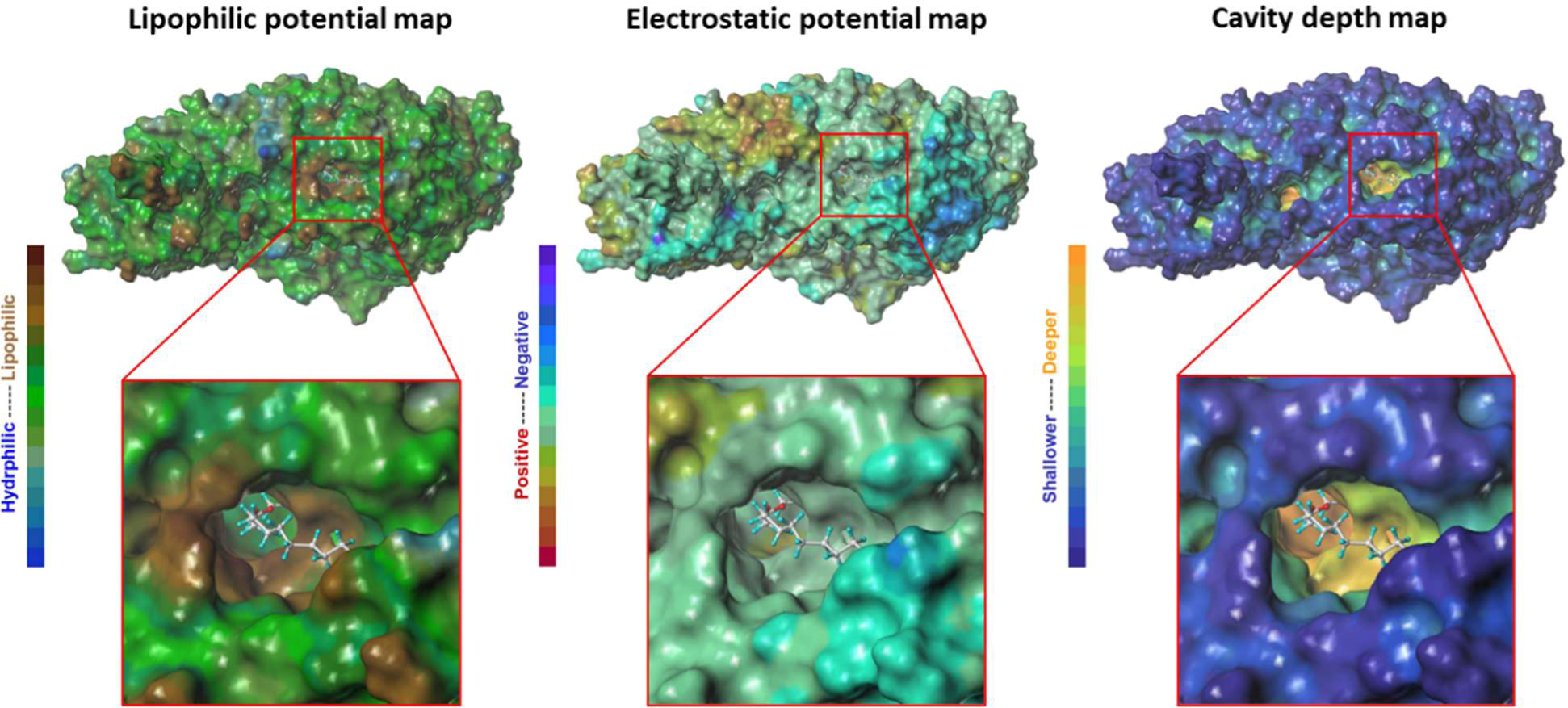
Lipophilic potential, electrostatic potential, and cavity depth maps generated for h15-LOX-2, with a close-up view of the entry to the arachidonic acid binding site. The maps were generated using the MOLCAD^91^ module of SYBYL-X^36^ v.2.1.1, applying the Connolly method^92^ for calculating the protein’s solvent accessible surface area. A substrate mimic inhibitor, which was co-crystallized with h15-LOX-2,^6^ is shown as ball-and-sticks (carbon, hydrogen, and oxygen atoms shown in gray, cyan, and red, respectively). The figure was prepared using SYBYL-X.

The proposed shape-based model was used to screen the “ZINC-Curated” database for compounds that have a high shape similarity to the model. ZINC-Curated is a “curated” version of the ZINC database,^33,34^ which was pre-filtered for potentially drug-like compounds, according to criteria defined in the “Materials and Methods” section.

Compounds from ZINC-Curated were ranked by ROCS^50^ according to their Shape Tanimoto^50^ score values. The 1,037 compounds with the highest Shape-Tanimoto values were selected for further analyses, which represents a reduction of ∼99.9% in the number of compounds from the ZINC-Curated database (**Figure 2A**). The Shape Tanimoto values for the selected compounds range from 0.516 to 0.565.

### 2D Structural “dissimilarity”^54^ filter

A preliminary visual inspection analysis of the compounds selected from the ZINC-Curated database by the shape-based model revealed that many had very similar structures among each other. Considering that we aim to select inhibitors with as diverse structures as possible, we applied a 2D structural “dissimilarity” filter in a second step of our VS (**Figure 2A**), aiming to eliminate compounds with high degrees of structural similarity to each other. Proceeding as described in the Materials and Methods section, structurally similar compounds were grouped into clusters based on molecular fingerprints.^54^ Using a “dissimilarity” index of 0.15 (*i.e*., a maximum of 85% of structural similarity) as a cut-off value, 514 clusters were generated. Only one representative compound of each cluster, randomly selected, was retained for the next steps, which means that 514 compounds were selected by the 2D structural “dissimilarity” filter. This represents a reduction of 50.4% in the number of compounds selected from the ZINC-Curated by our generated shape-based model (**Figure 2A**).

### Docking and visual inspection analyses

Compounds selected by the shape-based matching and by the 2D “dissimilarity” filters were subsequently docked into the h15-LOX-2 active site, using GOLD.^57^ The docking procedure was previously validated by “re-docking” the substrate mimic inhibitor C8E4, which was co-crystalized with h15-LOX-2 (PDB code: 4NRE),^6^ into the h15-LOX-2 active site. The re-docking was done using each of the four scoring functions available in GOLD^57^ (ChemPLP,^58^ ASP,^73^ Chemscore^57^ and Goldscore^74^). As shown in **Table S1**, the RMSD value calculated between Goldscore best-scored docking pose (pose 1) of C8E4 and the crystallographic pose of this inhibitor was the lowest (2.07 Å) compared to the other scoring functions. This value is lower than the resolution value of the complex 15-LOX-2/C8E4 crystal structure (2.63 Å). When considering the average of the two best-scored docking poses, ChemPLP also performs relatively well, with an average RMSD of 2.79 Å (**Table S1**). A visual inspection of the docking results confirms that the best-scored poses of both Goldscore and ChemPLP performed well in terms of reproducing the crystallographic binding pose of the inhibitor C8E4 (**Figure S1**). Therefore, only ChemPLP and Goldscore were used as scoring functions in the docking procedures of our VS protocol.

Among the 514 compounds submitted to docking into h15-LOX-2 active site’s cavity, the 50 with highest ChemPLP score values and the 50 with highest GoldScore score values were selected for subsequent analysis. As 10 were selected using both scoring functions, a total of 90 compounds were selected for the next step of our VS protocol, which represents a reduction of ∼82.5% in the number of compounds retained in the 2D “dissimilarity” filter step (**Figure 2A**).

The ChemPLP and Goldscore score values distributions of the compounds selected by the docking filter are shown in **Figure 4**. The ChemPLP score values range from 84 to 98, and the Goldscore score values range from 72 to 84. Notably, these score values are higher than the score values for the best-scored poses of the known substrate mimicking inhibitor C8E4^6^ (74.13 and 63.92 with ChemPLP and Goldscore, respectively), which was used for the docking validation procedure, as discussed above. To have an additional “positive control” of our docking procedures, the known inhibitor MLS000536924,^27^ which was used to build our shape-based model, was also submitted to docking into h15-LOX-2. Of note, the score values for this compound (64.36 and 59.90 with ChemPLP and Goldscore, respectively) were also lower compared to those of the compounds selected by our VS protocol (**Figure 4**).

**Figure 4.**
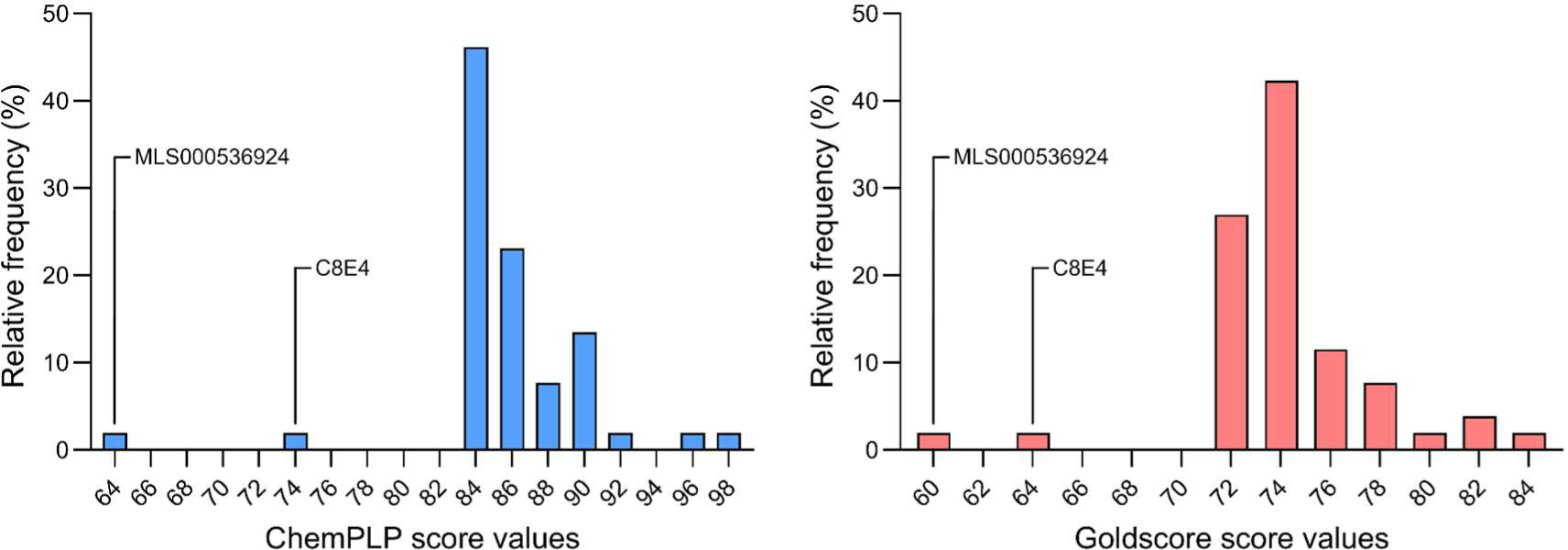
Score frequency distributions of the best-ranked docking poses for the compounds selected from the ZINC-Curated database after applying the docking filter. **(A)** ChemPLP score values distribution. **(B)** Goldscore score values distribution. The figure was prepared using GraphPad Prism. The ChemPLP and Goldscore score values of two h15-LOX-2 known inhibitors (MLS000536924^27^ and C8E4^6^) are represented for comparison.

In the last step of the proposed VS protocol, the best-scored pose of each compound selected by the docking filter was carefully analyzed by visual inspection inside the h15-LOX-2 active site’s cavity, using the LigandScout^53^ program for recognizing protein-ligand interactions. In this step, aiming to select the compounds that best fit into the h15-LOX-2 active site’s cavity, the following criteria were applied: (*i*) there should be hydrophobic interactions with at least four active site residues; (*ii*) there should be at least one hydrogen-bond and/or an ionic interaction and/or a π-cation interaction with at least one active site’s residue and/or with the catalytic iron and/or a water molecule; (*iii*) the best-scored docking poses should be “reproductible”, *i.e*., there should be at least four other poses, among the 10 docking poses, with similar conformations and orientations inside the active site’s cavity; and (*iv*) there should be an observable shape complementarity between the compound and the enzyme’s active site. Based on these criteria, 39 compounds were finally selected as h15-LOX-2 inhibitor candidates, which represents a reduction of ∼56.7% in the total of compounds selected by the docking filter (**Figure 2A**).

The structures of the selected compounds, as well as some of their physicochemical properties, are shown in **Table S2**. Among the 39 compounds selected from the ZINC-Curated database by our VS protocol, 14 (∼35.9%) could be acquired and tested in *in vitro* enzymatic assays for VS experimental validation (**Figure 2A**).

### Enzymatic inhibition assays for VS experimental validation and concluding discussion on the VS approach

To evaluate the inhibitory activities of the 14 acquired compounds (**1**-**14**, **Table 1**), *in vitro* inhibition assays with purified h15-LOX-2 were performed, using an UV-based method for detecting 15(*S*)-HpETE, the conjugate diene-containing enzyme-catalysed reaction product, at λ = 234 nm. h15-LOX-2 was expressed in *E. coli* BL21 DE3 pLysS cells and purified as described in the Materials and Methods section. A representative result of the SDS-PAGE analysis of h15-LOX-2 purified samples is shown in **Figure S3**. The values of the kinetic parameters K_M_, V_max_, and k_cat_, which were determined using arachidonic acid as a substrate, showed good agreement with the literature^6^ (**Figure S4** and **Table S3**). Dimethylsulfoxide (DMSO) was used as a solvent for the tested compounds, with a final concentration of 1.0% (v/v) in all assays. This DMSO concentration did not significantly affect h15-LOX-2 activity (**Figure S5**). All assays were performed in the presence of the non-ionic detergent Triton X-100 (at 0.01%, v/v) to avoid compound aggregate formation, which is recognized as an important source of false-positive results in enzyme inhibition screening assays, as discussed in the literature.^75–79^ At the concentration of 0.01% (v/v), Triton X-100 was relatively well tolerated by h15-LOX-2 (**Figure S6**).

**Table 1.**
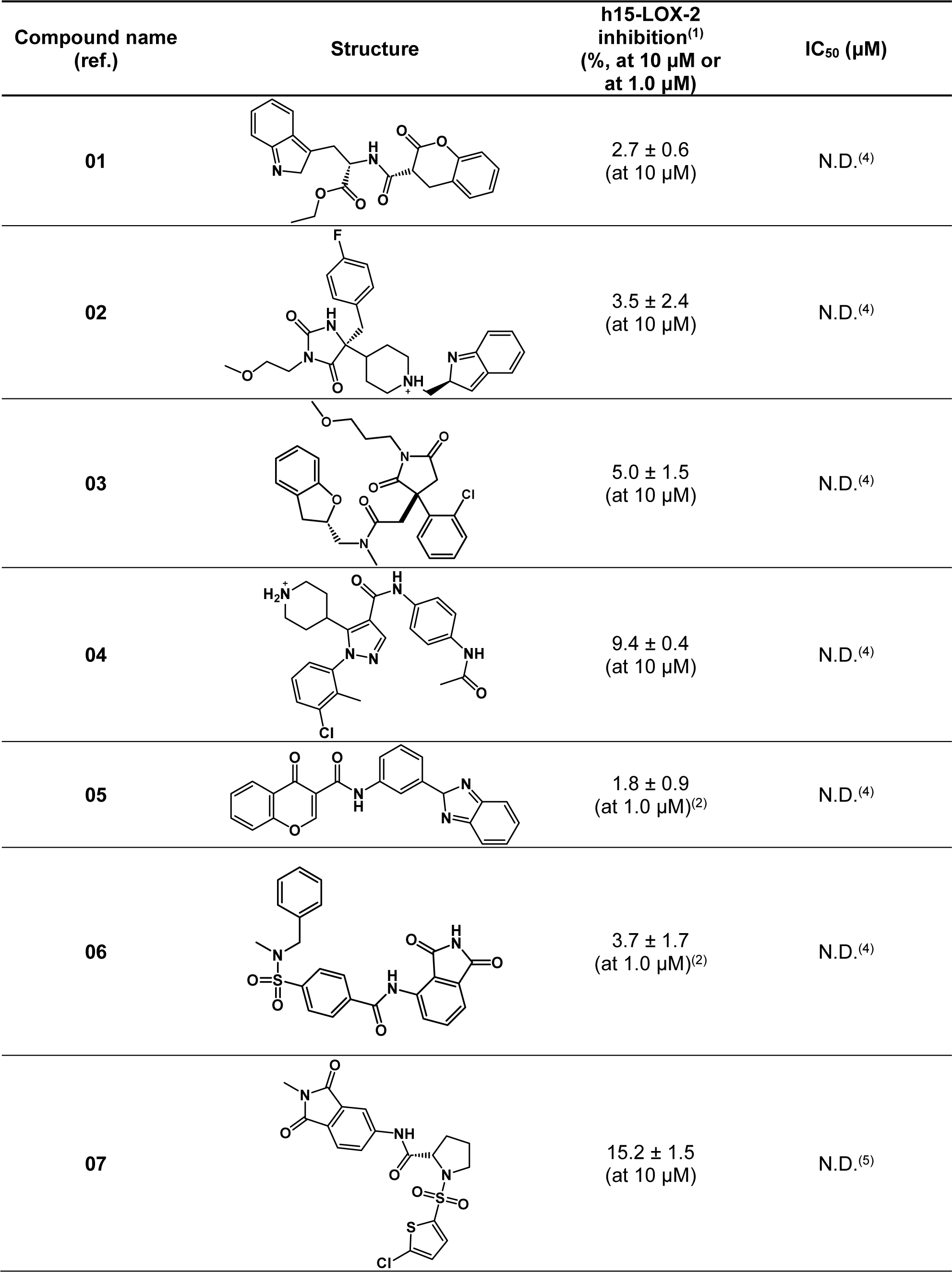
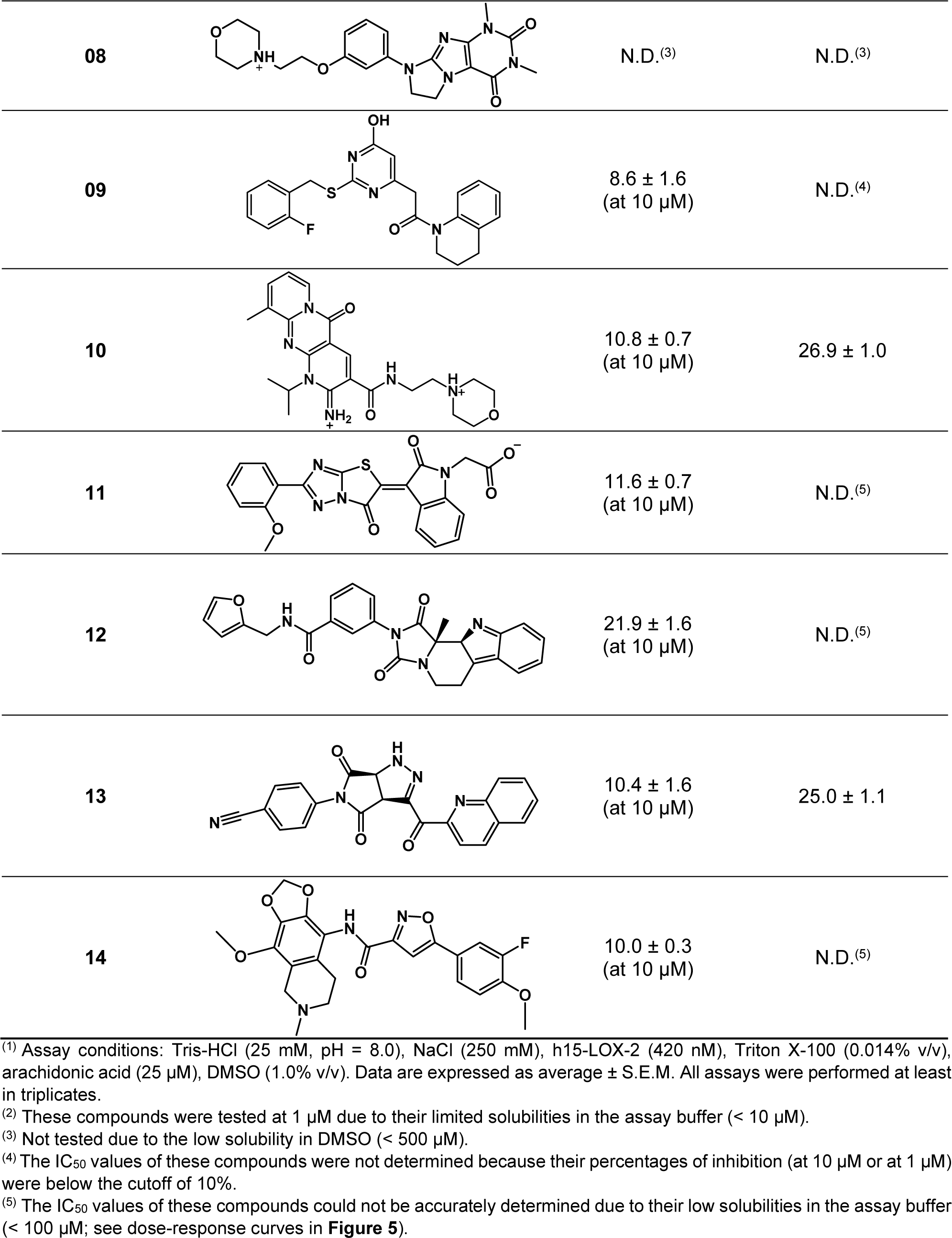
Structures and inhibitory activities (percentages of inhibition at 10 μM or at 1 μM) of the compounds selected as potential h15-LOX-2 inhibitors using the VS protocol proposed in this study.

| Compound name<br>(ref.) | Structure | h15-LOX-2<br>inhibition<sup>(1)</sup><br>(%, at 10 $\mu\text{M}$ or<br>at 1.0 $\mu\text{M}$) | IC<sub>50</sub> ($\mu\text{M}$) |
| --- | --- | --- | --- |
| 01 | | 2.7 ± 0.6<br>(at 10 $\mu\text{M}$) | N.D.<sup>(4)</sup> |
| 02 | | 3.5 ± 2.4<br>(at 10 $\mu\text{M}$) | N.D.<sup>(4)</sup> |
| 03 | | 5.0 ± 1.5<br>(at 10 $\mu\text{M}$) | N.D.<sup>(4)</sup> |
| 04 | | 9.4 ± 0.4<br>(at 10 $\mu\text{M}$) | N.D.<sup>(4)</sup> |
| 05 | | 1.8 ± 0.9<br>(at 1.0 $\mu\text{M}$)<sup>(2)</sup> | N.D.<sup>(4)</sup> |
| 06 | | 3.7 ± 1.7<br>(at 1.0 $\mu\text{M}$)<sup>(2)</sup> | N.D.<sup>(4)</sup> |
| 07 | | 15.2 ± 1.5<br>(at 10 $\mu\text{M}$) | N.D.<sup>(5)</sup> |

**Table 1. (Continued)**
|  |  |  |  |
| --- | --- | --- | --- |
| 08 |  | N.D.<sup>(3)</sup> | N.D.<sup>(3)</sup> |
| 09 |  | 8.6 ± 1.6<br>(at 10 μM) | N.D.<sup>(4)</sup> |
| 10 |  | 10.8 ± 0.7<br>(at 10 μM) | 26.9 ± 1.0 |
| 11 |  | 11.6 ± 0.7<br>(at 10 μM) | N.D.<sup>(5)</sup> |
| 12 |  | 21.9 ± 1.6<br>(at 10 μM) | N.D.<sup>(5)</sup> |
| 13 |  | 10.4 ± 1.6<br>(at 10 μM) | 25.0 ± 1.1 |
| 14 |  | 10.0 ± 0.3<br>(at 10 μM) | N.D.<sup>(5)</sup> |

<sup>(1)</sup> Assay conditions: Tris-HCl (25 mM, pH = 8.0), NaCl (250 mM), h15-LOX-2 (420 nM), Triton X-100 (0.014% v/v), arachidonic acid (25 μM), DMSO (1.0% v/v). Data are expressed as average ± S.E.M. All assays were performed at least in triplicates.
<sup>(2)</sup> These compounds were tested at 1 μM due to their limited solubilities in the assay buffer (< 10 μM).
<sup>(3)</sup> Not tested due to the low solubility in DMSO (< 500 μM).
<sup>(4)</sup> The IC<sub>50</sub> values of these compounds were not determined because their percentages of inhibition (at 10 μM or at 1 μM) were below the cutoff of 10%.
<sup>(5)</sup> The IC<sub>50</sub> values of these compounds could not be accurately determined due to their low solubilities in the assay buffer (< 100 μM; see dose-response curves in **Figure 5**).

Initially, all 14 compounds were tentatively tested at 10 μM (or at 1 μM, depending on the solubility in the assay buffer) (**Table 1**). Compound **08** could not be tested due to its low solubility in DMSO (< 500 μM). Dose-response curves were obtained for five compounds that showed at least 10% of inhibition at 10 μM: compounds **07**, **10**, **11**, **12**, and **13** (**Figure 5**). Compounds **10** and **13** showed IC_50_ values of (26.9 ± 1.0) μM and (25.0 ± 1.1) μM, respectively (**Table 1**). Due to the limited solubilities of **07**, **11**, and **12** in the assay buffer (< 100 μM), only incomplete dose-response curves were obtained for these compounds (**Figure 5**) and their IC_50_ values could not be accurately determined. A dose-response curve could not be obtained for **14**, despite this compound showed more than 10% of inhibition at 10 μM, because it was not soluble in the assay buffer at concentrations higher than 50 μM. Nonetheless, it was possible to observe that **07** and **11** showed more than 50% of inhibition at 25.1 μM and at 63.1 μM, respectively.

**Figure 5.**
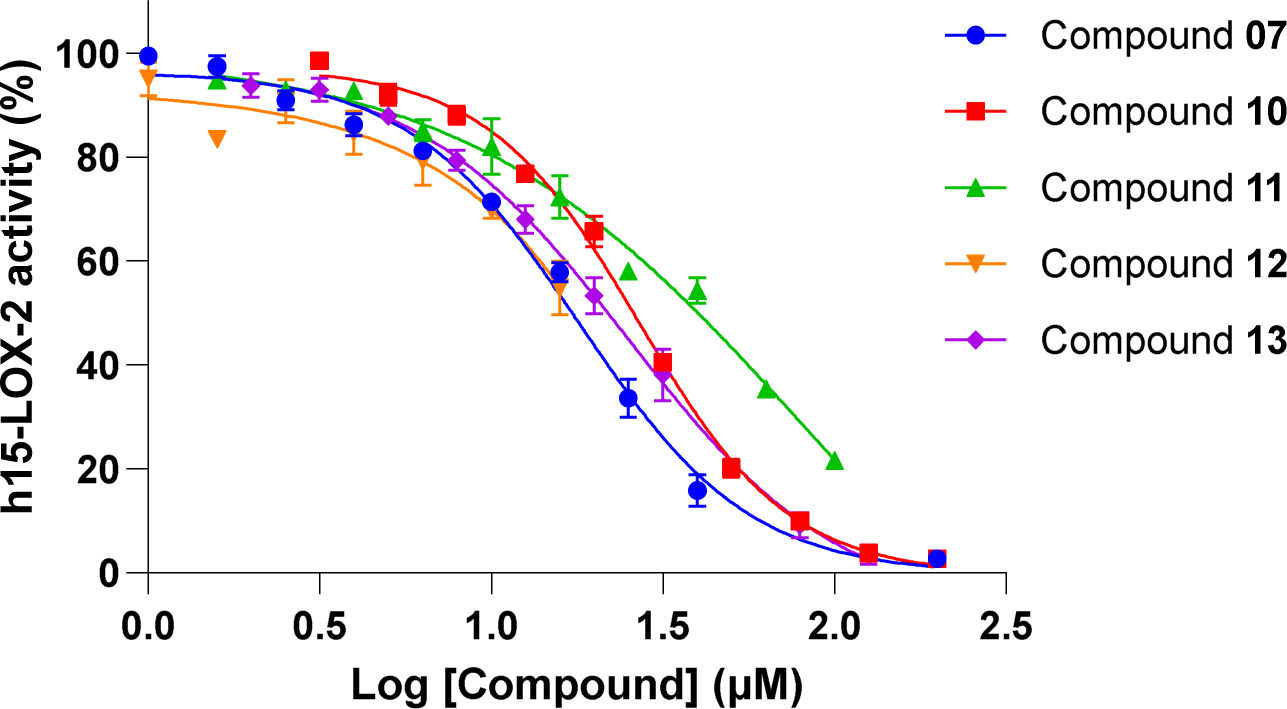
Dose-response curves for compounds 07, 10, 11, 12, and 13. All assays were carried out in a total volume of 300 μL, containing Tris-HCl (25 mM, pH = 8.0), NaCl (250 mM), h15-LOX-2 (420 nM), Triton X-100 (0.014% v/v), arachidonic acid (25 μM), and each tested compound (1.0 μM to up to 199.1 μM, depending on the compound solubility in the assay buffer). Data represent the average ± S.E.M. of experiments performed at least in triplicates. The assays with compounds **10** and **13** were performed twice. Complete dose-response curves could not be obtained for compounds **07**, **11**, and **12**, due to their low solubilities in the assay buffer (< 100 μM). The figure was prepared using GraphPad Prism.

The low solubilities of some inhibitors selected by our VS are expected, even though we used solubility filters based on *in silico* predictions^38^ in the preparation of the ZINC-Curated database (see Materials and Methods section). First, as discussed in the literature,^63^ the lipophilic nature of the substrate binding pockets in fatty acid-binding proteins, such as the h15-LOX-2 active site’s cavity (see **Figure 2A**), poses a considerable challenge in terms of design of soluble and “drug-like” inhibitors. Second, despite the advances in the field, accurate *in silico* prediction of solubility remains a big challenge due to the complexity of the chemical and physicochemical factors that influence solubility,^38,80–82^ including the nature, temperature, pH, and ionic strength of the solvent, solid state of the solute (amorph or crystal), solute polymorphisms, intermolecular interactions between the solute and the solvent, ionization state(s) of the solute, among others. Despite the solubility issues, compounds **07**, **11**, **12**, and **14**, as well as the other compounds selected by our VS and tested through enzymatic assays, passed the “drug-like” filter criteria that were used for building our ZINC-Curated database, in addition to having structural and physicochemical properties that are overall inside the suitable physicochemical space for oral bioavailability, according to Lipinki’s Rule of Five^41,42^ and other empirical rules, such as those implemented in SWISS ADME^83^ (**Figure S2**).

Lineweaver-Burk plots revealed a mixed-type inhibition for either **10** and **13** (**Figure 7**), with K_i_ values of (16.4 ± 8.1) μM and (15.1 ± 7.6) μM, respectively, and α values of (1.4 ± 1.1) and (4.3 ± 2.8), respectively. As discussed in the literature,^84–86^ there are several mechanisms of protein-inhibitor binding that would be compatible with a mixed-type inhibition model, which is also treated as a special case of “noncompetitive” inhibition by some authors.^84–86^ According to the conventional “two-site” model, mixed-type inhibitors would bind both the free enzyme and the enzyme-substrate complex.^84–86^ However, recent studies and reviews from the literature have been suggested that in most cases the molecular nature of mixed-type inhibition is mainly competitive, which means that mixed-type inhibitors would primarily bind the free enzyme, and only at the substrate binding site.^84–86^ Therefore, considering that our VS aimed at selecting compounds that bind the h15-LOX-2 active site’s cavity, where the substrate binds, the experimentally-evidenced mixed-type inhibition models of **10** and **13** are compatible with the predicted binding modes of these two inhibitors to h15-LOX-2.

**Figure 6.**
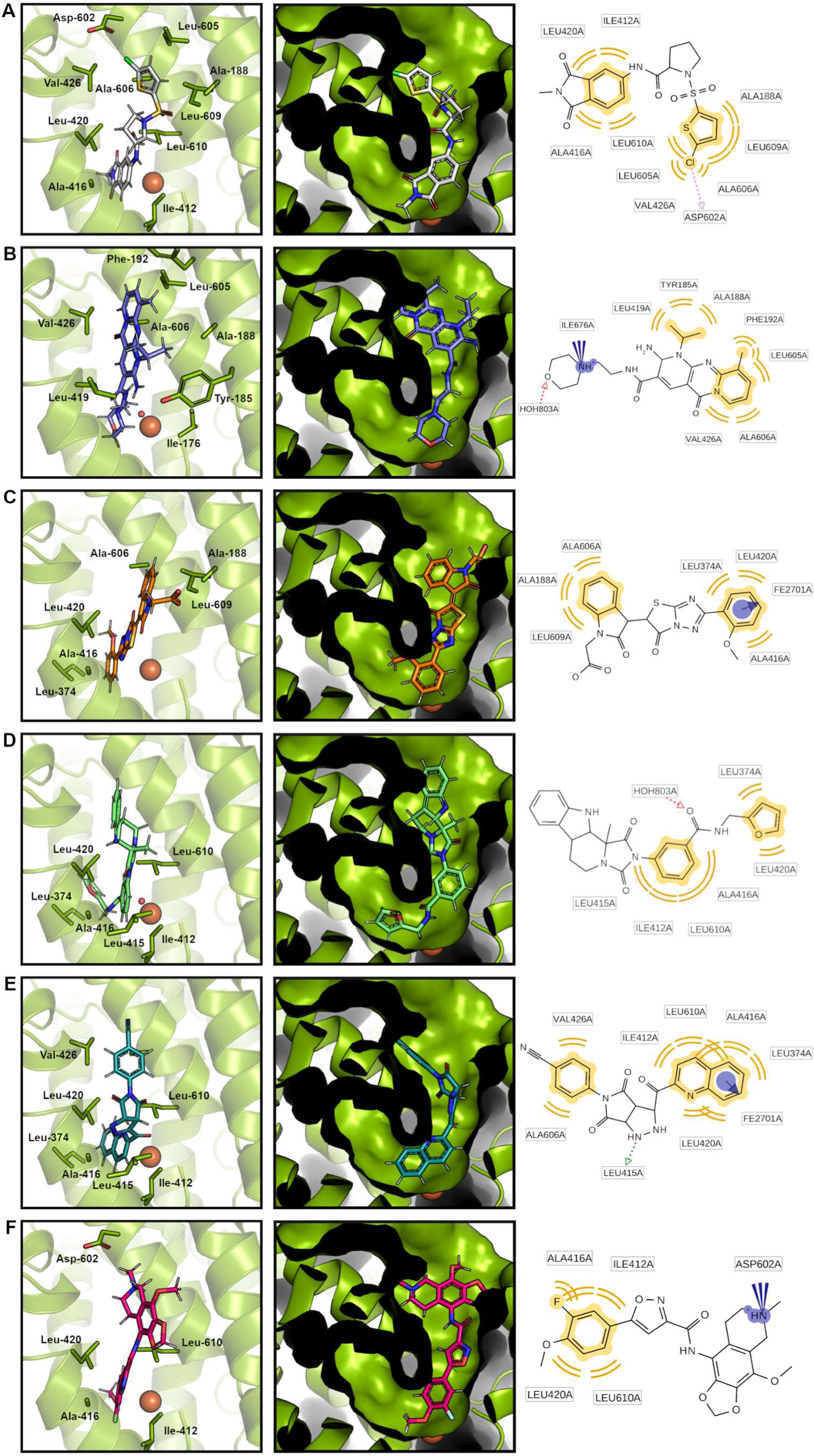
Schematic representations of the predicted binding modes (best-scored docking solutions) of the h15-LOX-2 inhibitors selected by the VS protocol proposed in this study: (A) Compound 07; (B) Compound 10; (C) Compound 11; (D) Compound 12; (E) Compound 13; and (F) Compound 14. *Left panels*: Schematic representations of the best-scored docking solutions for each inhibitor inside the h15-LOX-2 active site’s cavity. For clarity, only the active site’s residues that interact with each inhibitor are highlighted (shown as sticks). Catalytic iron is shown as a brown sphere. A water molecule that coordinates iron and that makes a hydrogen-bond interaction with compounds **10** and **12** is shown as a red sphere in panels (B) and (D). *Central panels*: Schematic representations of the best-scored docking solutions for each compound, with h15-LOX-2 active site’s cavity represented as a surface to highlight its U-shaped format. *Right panels*: 2D Schematic representations of the protein-ligand interactions identified using the LigandScout^53^ program. Yellow: hydrophobic interactions; blue circle/arrow: pi-cation interactions; blue cones: ionic interactions; red arrows: hydrogen-bond acceptor interactions. The figure was prepared using PyMOL.

**Figure 7.**
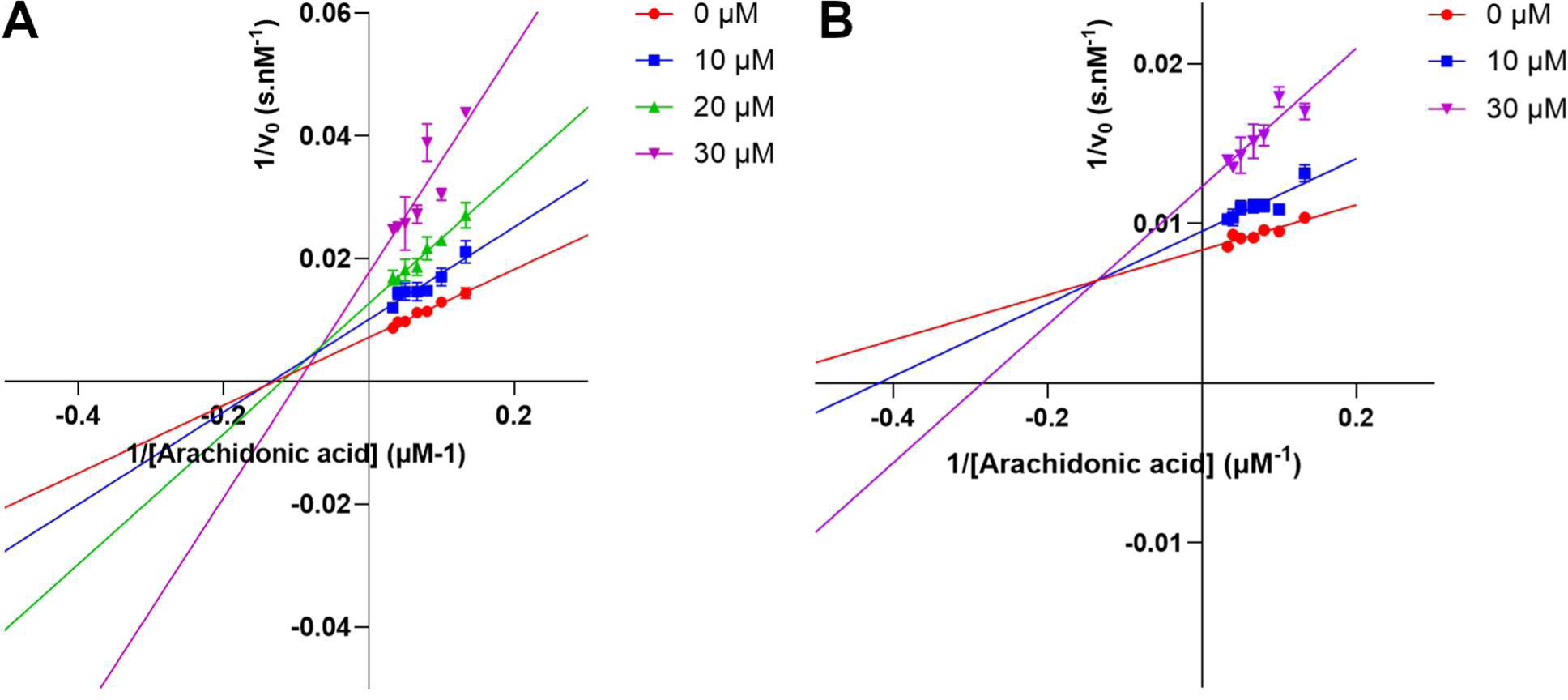
Lineweaver-Burk plots for: (A) Compound 10 and (B) Compound 13. All assays were carried out in a total volume of 300 μL, containing Tris-HCl (25 mM, pH = 8.0), NaCl (250 mM), h15-LOX-2 (420 nM), Triton X-100 (0.014% v/v), arachidonic acid (7.5 μM - 30 μM), and the tested compound (10 μM, 20 μM, and/or 30 μM). Data represent the average ± S.E.M. of experiments performed at least in triplicates. The figure was prepared using GraphPad Prism.

According to docking predictions, all six inhibitors that showed inhibitory activities higher than 10% at 10 μM interact with h15-LOX-2 mainly *via* hydrophobic interactions with the residues that form the enzyme’s active site cavity (**Figure 6**). This is expected, considering that this cavity is highly lipophilic (as shown in **Figure 3**) and formed by the side chains of several hydrophobic amino acids, such as Ala-188, Leu-374, Ile-412, Leu-415, Leu-420, Ala-416, Val-426, Leu-605, Ala-606, Leu-609, and Leu-610 (**Figure 1**), which accommodate the lipid substrate arachidonic acid. The identified inhibitors are predicted to occupy the more solvent-exposed arm of the U-shaped active site’s cavity (**Figure 6**, central panels), and possibly block the substrate access to the entrance of this cavity. Notably, π-cation interactions are predicted to occur between h15-LOX-2 catalytic iron and the aromatic systems of **11** and of **13** (**Figure 6**). An ionic interaction is expected between the charged nitrogen of **10** morpholine ring and the carboxylate group of C-terminal Ile-676, and a hydrogen-bond could be established between the oxygen of the ring and an iron coordinating water molecule (**Figure 6**). Shape complementarity between the identified inhibitors and their binding sites inside the active site’s cavity was clearly observed in the visual inspection analysis, which reinforces the effectiveness of our shape-based approach used as a first filter in the proposed VS.

One should notice that the identified inhibitors have diverse structures from each other and from the h15-LOX-2 inhibitors reported in the literature,^8,27,28^ which maximizes the number of scaffolds that can be used as starting points for a hit-to-lead optimization project.^87^ The achieved structural diversity is highly desirable in a VS campaign,^87^ and it is particularly relevant in the search of h15-LOX-2 inhibitors, considering that h15-LOX-2 is a very underexploited drug target, and there is a small number of h15-LOX-2 inhibitors described in the literature so far, with most of them deriving from similar scaffolds.^8,27,28^ The success in the selection of structurally diverse inhibitors in our VS is related to the choice of selection filters we used. As presented, a shape-based approach was used as the first selection filter. It is known that a major advantage of a shape-based approach is that it allows for selecting compounds having different scaffolds, despite sharing a similar shape.^50,72,88,89^ We additionally improved the structural diversity of the dataset by using a 2D structural “dissimilarity” approach to group similar compounds into clusters, aiming to eliminate compounds that had more than 85% of similarity among each other.

## CONCLUSIONS

In summary, in the VS presented here, we successfully combined ligand-based approaches^32^ (shape-based matching and a 2D “dissimilarity” filter) and a structure-based approach^32^ (docking followed by careful visual inspection) to screen a “curated” version of the ZINC database, pre-filtered for drug-like compounds (here called “ZINC-Curated”), for novel h15-LOX-2 inhibitors. Two inhibitors, **10** and **13**, with K_i_ values of (16.4 ± 8.1) μM and (15.1 ± 7.6) μM, respectively, were identified, revealing a mixed-type mechanism of inhibition. Four additional compounds (**07**, **11**, **12**, and **14**) were found to inhibit h15-LOX-2 in the micromolar range, but the IC_50_ and K_i_ values of these compounds could not be accurately calculated due to their low solubilities in the enzymatic assays buffer. The inhibitors identified in this study are structurally diverse from each other and from the h15-LOX-2 inhibitors reported in the literature,^8,27,28^ and are expected to have good oral bioavailabilities based on empirical rules.^41,42,90^ Binding modes predicted by docking reveal that, not surprisingly, all identified inhibitors interact with the h15-LOX-2 active site’s cavity residues mainly by hydrophobic interactions, and they possibly occupy the more solvent-exposed arm of the U-shaped enzyme’s active site cavity, blocking the entrance of arachidonic acid to this cavity. Despite an in-depth characterization of the binding modes of the identified inhibitors through experimental techniques are outside the scope of this study, their inhibitory potencies make them useful as starting points for hit-to-lead optimization projects aiming at improving bind affinities and, eventually, selectivities and pharmacokinetic properties. From a biological perspective, *in vitro* assays for evaluating the effects of the identified inhibitors in lipid accumulation, h15-LOX-2-derived lipid mediators’ distribution, and foam cell formation in cell-based atherosclerosis models might be considered as a further step. Studies in this direction are underway.

Finally, it is important to stress that, to our knowledge, this study was the first to exploit a shape-based approach to search for novel h15-LOX-2 inhibitors. Considering the importance of shape complementarity for protein-ligand recognition, specially inside the lipophilic and buried binding sites of proteins that bind lipids, such as h15-LOX-2, the choice for a shape-based approach as a first filter in our VS protocol has been crucial for the VS success. Given the increasing evidence on the possible role of h15-LOX-2 in atherosclerosis pathogenesis,^9–15^ as well as in other diseases and pathological processes,^8,17,21,23^ including inflammation and, possibly, ferroptosis, the here reported identification of novel inhibitors of this so-far underexploited target enzyme is of major importance.

## AUTHOR INFORMATION

### Notes

The authors declare no competing financial interest.

## ACKNOWLEDGMENTS

L.G.V. acknowledges his PhD fellowship from FAPESP (2021/10514-8); T.S.I. thanks her MSc fellowship from CAPES (88887.603627/2021-00); L.E.S.N., A.T.-do A. and S.M. acknowledge CEPID Redoxoma (2013/07937-8) for financial support.

We acknowledge Prof. Dr. Marcia Newcomer (Louisiana State University, USA) for kindly providing the plasmid for h15-LOX-2 expression and Prof. Dr. Claus Schneider (Vanderbilt University, USA) for his valuable scientific discussion. We also thank Prof. Dr. Sandro Roberto Marana, Prof. Dr. Ohara Augusto, and Fernando Rodrigues Coelho (University of São Paulo) for their assistance in the production of recombinant h15-LOX-2. Additionally, we acknowledge Openeye Scientific Sofware Inc. (Santa Fe, USA) for ROCS, OMEGA, and QUACPAC software licenses, and Prof. Dr. Thierry Langer (Inte:Ligand GmbH and University of Vienna, Vienna, Austria) for kindly providing the LigandScout software license.

## REFERENCES

(1) Kuhn, H.; Banthiya, S.; van Leyen, K. Mammalian Lipoxygenases and Their Biological Relevance. Biochimica et Biophysica Acta - Molecular and Cell Biology of Lipids. Elsevier B.V. 2015, pp 308–330. 10.1016/j.bbalip.2014.10.002.

(2) Schneider, C.; Pratt, D. A.; Porter, N. A.; Brash, A. R. Control of Oxygenation in Lipoxygenase and Cyclooxygenase Catalysis. Chemistry and Biology. May 29, 2007, pp 473–488. 10.1016/j.chembiol.2007.04.007.

(3) Kuhn, H.; Humeniuk, L.; Kozlov, N.; Roigas, S.; Adel, S.; Heydeck, D. The Evolutionary Hypothesis of Reaction Specificity of Mammalian ALOX15 Orthologs. Progress in Lipid Research. Elsevier Ltd October 1, 2018, pp 55–74. 10.1016/j.plipres.2018.09.002.

(4) Oliw, E. H. Thirty Years with Three-Dimensional Structures of Lipoxygenases. Archives of Biochemistry and Biophysics. Academic Press Inc. February 1, 2024. 10.1016/j.abb.2023.109874.

(5) Dyall, S. C.; Balas, L.; Bazan, N. G.; Brenna, J. T.; Chiang, N.; da Costa Souza, F.; Dalli, J.; Durand, T.; Galano, J. M.; Lein, P. J.; Serhan, C. N.; Taha, A. Y. Polyunsaturated Fatty Acids and Fatty Acid-Derived Lipid Mediators: Recent Advances in the Understanding of Their Biosynthesis, Structures, and Functions. Progress in Lipid Research. Elsevier Ltd April 1, 2022. 10.1016/j.plipres.2022.101165.

(6) Kobe, M. J.; Neau, D. B.; Mitchell, C. E.; Bartlett, S. G.; Newcomer, M. E. The Structure of Human 15-Lipoxygenase-2 with a Substrate Mimic. Journal of Biological Chemistry 2014, 289 (12), 8562–8569. 10.1074/jbc.M113.543777.

(7) Mashima, R.; Okuyama, T. The Role of Lipoxygenases in Pathophysiology; New Insights and Future Perspectives. Redox Biology. Elsevier B.V. December 1, 2015, pp 297–310. 10.1016/j.redox.2015.08.006.

(8) Tsai, W. C.; Gilbert, N. C.; Ohler, A.; Armstrong, M.; Perry, S.; Kalyanaraman, C.; Yasgar, A.; Rai, G.; Simeonov, A.; Jadhav, A.; Standley, M.; Lee, H. W.; Crews, P.; Iavarone, A. T.; Jacobson, M. P.; Neau, D. B.; Offenbacher, A. R.; Newcomer, M.; Holman, T. R. Kinetic and Structural Investigations of Novel Inhibitors of Human Epithelial 15-Lipoxygenase-2. Bioorg Med Chem 2021, 46. 10.1016/j.bmc.2021.116349.

(9) Snodgrass, R. G.; Brüne, B. Regulation and Functions of 15-Lipoxygenases in Human Macrophages. Frontiers in Pharmacology. Frontiers Media S.A. July 4, 2019. 10.3389/fphar.2019.00719.

(10) Benatzy, Y.; Palmer, M. A.; Lütjohann, D.; Ohno, R. I.; Kampschulte, N.; Schebb, N. H.; Fuhrmann, D. C.; Snodgrass, R. G.; Brüne, B. ALOX15B Controls Macrophage Cholesterol Homeostasis via Lipid Peroxidation, ERK1/2 and SREBP2. Redox Biol 2024, 72. 10.1016/j.redox.2024.103149.

(11) Rydberg, E. K.; Krettek, A.; Ullström, C.; Ekström, K.; Svensson, P. A.; Carlsson, L. M. S.; Jönsson-Rylander, A. C.; Hansson, G. I.; McPheat, W.; Wiklund, O.; Ohlsson, B. G.; Hultén, L. M. Hypoxia Increases LDL Oxidation and Expression of 15-Lipoxygenase-2 in Human Macrophages. Arterioscler Thromb Vasc Biol 2004, 24 (11), 2040–2045. 10.1161/01.ATV.0000144951.08072.0b.

(12) Gertow, K.; Nobili, E.; Folkersen, L.; Newman, J. W.; Pedersen, T. L.; Ekstrand, J.; Swedenborg, J.; Kühn, H.; Wheelock, C. E.; Hansson, G. K.; Hedin, U.; Haeggström, J. Z.; Gabrielsen, A. 12- and 15-Lipoxygenases in Human Carotid Atherosclerotic Lesions: Associations with Cerebrovascular Symptoms. Atherosclerosis 2011, 215 (2), 411–416. 10.1016/j.atherosclerosis.2011.01.015.

(13) Hultén, L. M.; Olson, F. J.; Åberg, H.; Carlsson, J.; Karlström, L.; Borén, J.; Fagerberg, B.; Wiklund, O. 15-Lipoxygenase-2 Is Expressed in Macrophages in Human Carotid Plaques and Regulated by Hypoxia-Inducible Factor-1α. Eur J Clin Invest 2010, 40 (1), 11–17. 10.1111/j.1365-2362.2009.02223.x.

(14) Magnusson, L. U.; Lundqvist, A.; Karlsson, M. N.; Skålén, K.; Levin, M.; Wiklund, O.; Borén, J.; Hultén, L. M. Arachidonate 15-Lipoxygenase Type B Knockdown Leads to Reduced Lipid Accumulation and Inflammation in Atherosclerosis. PLoS One 2012, 7 (8). 10.1371/journal.pone.0043142.

(15) Danielsson, K. N.; Rydberg, E. K.; Ingelsten, M.; Akyürek, L. M.; Jirholt, P.; Ullström, C.; Forsberg, G. B.; Borén, J.; Wiklund, O.; Hultén, L. M. 15-Lipoxygenase-2 Expression in Human Macrophages Induces Chemokine Secretion and T Cell Migration. Atherosclerosis 2008, 199 (1), 34–40. 10.1016/j.atherosclerosis.2007.10.027.

(16) Barooni, A. B.; Ghorbani, M.; Salimi, V.; Alimohammadi, A.; Khamseh, M. E.; Akbari, H.; Imani, M.; Nourbakhsh, M.; Sheikhi, A.; Shirian, F. I.; Ameri, M.; Tavakoli-Yaraki, M. Up-Regulation of 15-Lipoxygenase Enzymes and Products in Functional and Non-Functional Pituitary Adenomas. Lipids Health Dis 2019, 18 (1). 10.1186/s12944-019-1089-1.

(17) Yang, L.; Ma, C.; Zhang, L.; Zhang, M.; Li, F.; Zhang, C.; Yu, X.; Wang, X.; He, S.; Zhu, D.; Song, Y. 15-Lipoxygenase-2/15(S)-Hydroxyeicosatetraenoic Acid Regulates Cell Proliferation and Metastasis via the STAT3 Pathway in Lung Adenocarcinoma. Prostaglandins Other Lipid Mediat 2018, 138, 31–40. 10.1016/j.prostaglandins.2018.07.003.

(18) Wang, D.; Chen, S.; Feng, Y.; Yang, Q.; Campbell, B. H.; Tang, X.; Campbell, W. B. Reduced Expression of 15-Lipoxygenase 2 in Human Head and Neck Carcinomas. Tumor Biology 2006, 27 (5), 261–273. 10.1159/000094761.

(19) Jiang, W. G.; Watkins, G.; Douglas-Jones, A.; Mansel, R. E. Reduction of Isoforms of 15-Lipoxygenase (15-LOX)-1 and 15-LOX-2 in Human Breast Cancer. Prostaglandins Leukot Essent Fatty Acids 2006, 74 (4), 235–245. 10.1016/j.plefa.2006.01.009.

(20) Feng, Y.; Bai, X.; Yang, Q.; Wu, H.; Wang, D. Downregulation of 15-Lipoxygenase 2 by Glucocorticoid Receptor in Prostate Cancer Cells. Int J Oncol 2010, 36 (6), 1541–1549. 10.3892/ijo_00000641.

(21) Subbarayan, V.; Xu, X. C.; Kim, J.; Yang, P.; Hoque, A.; Sabichi, A. L.; Llansa, N.; Mendoza, G.; Logothetis, C. J.; Newman, R. A.; Lippman, S. M.; Menter, D. G. Inverse Relationship between 15-Lipoxygenase-2 and PPAR-γ Gene Expression in Normal Epithelia Compared with Tumor Epithelia. Neoplasia 2005, 7 (3), 280–293. 10.1593/neo.04457.

(22) Schneider, C.; Pozzi, A. Cyclooxygenases and Lipoxygenases in Cancer. Cancer and Metastasis Reviews. December 2011, pp 277–294. 10.1007/s10555-011-9310-3.

(23) Daurkin, I.; Eruslanov, E.; Stoffs, T.; Perrin, G. Q.; Algood, C.; Gilbert, S. M.; Rosser, C. J.; Su, L. M.; Vieweg, J.; Kusmartsev, S. Tumor-Associated Macrophages Mediate Immunosuppression in the Renal Cancer Microenvironment by Activating the 15-Lipoxygenase-2 Pathway. Cancer Res 2011, 71 (20), 6400–6409. 10.1158/0008-5472.CAN-11-1261.

(24) Pratt, D. A. Targeting Lipoxygenases to Suppress Ferroptotic Cell Death. Proceedings of the National Academy of Sciences of the United States of America. National Academy of Sciences 2023. 10.1073/PNAS.2309317120.

(25) Conrad, M.; Pratt, D. A. The Chemical Basis of Ferroptosis. Nat Chem Biol 2019, 15 (12), 1137–1147. 10.1038/s41589-019-0408-1.

(26) Shah, R.; Shchepinov, M. S.; Pratt, D. A. Resolving the Role of Lipoxygenases in the Initiation and Execution of Ferroptosis. ACS Cent Sci 2018, 4 (3), 387–396. 10.1021/acscentsci.7b00589.

(27) Jameson, J. B.; Kantz, A.; Schultz, L.; Kalyanaraman, C.; Jacobson, M. P.; Maloney, D. J.; Jadhav, A.; Simeonov, A.; Holman, T. R. A High Throughput Screen Identifies Potent and Selective Inhibitors to Human Epithelial 15-Lipoxygenase-2. PLoS One 2014, 9 (8). 10.1371/journal.pone.0104094.

(28) Vasquez-Martinez, Y.; Ohri, R. V.; Kenyon, V.; Holman, T. R.; Sepúlveda-Boza, S. Structure-Activity Relationship Studies of Flavonoids as Potent Inhibitors of Human Platelet 12-HLO, Reticulocyte 15-HLO-1, and Prostate Epithelial 15-HLO-2. Bioorg Med Chem 2007, 15 (23), 7408–7425. 10.1016/j.bmc.2007.07.036.

(29) Dar, H. H.; Mikulska-Ruminska, K.; Tyurina, Y. Y.; Luci, D. K.; Yasgar, A.; Samovich, S. N.; Kapralov, A. A.; Souryavong, A. B.; Tyurin, V. A.; Amoscato, A. A.; Epperly, M. W.; Shurin, G. V.; Standley, M.; Holman, T. R.; Croix, C. M. S.; Watkins, S. C.; VanDemark, A. P.; Rana, S.; Zakharov, A. V.; Simeonov, A.; Marugan, J.; Mallampalli, R. K.; Wenzel, S. E.; Greenberger, J. S.; Rai, G.; Bayir, H.; Bahar, I.; Kagan, V. E. Discovering Selective Antiferroptotic Inhibitors of the 15LOX/PEBP1 Complex Noninterfering with Biosynthesis of Lipid Mediators. Proc Natl Acad Sci U S A 2023, 120 (25). 10.1073/pnas.2218896120.

(30) Osman, N. A.; Soltan, M. K.; Rezq, S.; Flaherty, J.; Romero, D. G.; Abdelkhalek, A. S. Dual COX-2 and 15-LOX Inhibition Study of Novel 4-Arylidine-2-Mercapto-1-Phenyl-1H-Imidazolidin-5(4H)-Ones: Design, Synthesis, Docking, and Anti-Inflammatory Activity. Arch Pharm (Weinheim) 2024, 357 (5). 10.1002/ardp.202300615.

(31) Tran, M.; Yang, K.; Glukhova, A.; Holinstat, M.; Holman, T. Inhibitory Investigations of Acyl-CoA Derivatives against Human Lipoxygenase Isozymes. Int J Mol Sci 2023, 24 (13). 10.3390/ijms241310941.

(32) Klebe, G. Virtual Ligand Screening: Strategies, Perspectives and Limitations. Drug Discov Today 2006, 11 (13–14), 580–594. 10.1016/j.drudis.2006.05.012.

(33) Irwin, J. J.; Sterling, T.; Mysinger, M. M.; Bolstad, E. S.; Coleman, R. G. ZINC: A Free Tool to Discover Chemistry for Biology. Journal of Chemical Information and Modeling. American Chemical Society July 23, 2012, pp 1757–1768. 10.1021/ci3001277.

(34) Irwin, J. J.; Shoichet, B. K. ZINC − A Free Database of Commercially Available Compounds for Virtual Screening ZINC - A Free Database of Commercially Available Compounds for Virtual Screening. J. Chem. Inf. Model 2005, 45 (December 2004), 177–182. 10.1021/ci049714.

(35) Leeson, P. D.; Bento, A. P.; Gaulton, A.; Hersey, A.; Manners, E. J.; Radoux, C. J.; Leach, A. R. Target-Based Evaluation of “Drug-Like” Properties and Ligand Efficiencies. Journal of Medicinal Chemistry. American Chemical Society June 10, 2021, pp 7210–7230. 10.1021/acs.jmedchem.1c00416.

(36) SYBYL. SYBYL-X. Tripos, a Certara Companie: 1699 South Hanley Rd., St. Louis, Missouri, 63144, USA 2013.

(37) Powell, M. J. D. Restart Procedures for the Conjugate Gradient Method. Math Program 1977, 12, 241–254.

(38) Hill, A. P.; Young, R. J. Getting Physical in Drug Discovery: A Contemporary Perspective on Solubility and Hydrophobicity. Drug Discov Today 2010, 15 (15–16), 648–655. 10.1016/j.drudis.2010.05.016.

(39) Oprea, T. I. Property Distribution of Drug-Related Chemical Databases. J Comput Aided Mol Des 2000, 14 (3), 251–264. 10.1023/A:1008130001697.

(40) FILTER v.2.1.1. OpenEye Scientific Software: Santa Fe, NM, USA.

(41) Lipinski, C. A. Rule of Five in 2015 and beyond: Target and Ligand Structural Limitations, Ligand Chemistry Structure and Drug Discovery Project Decisions. Advanced Drug Delivery Reviews. Elsevier B.V. June 1, 2016, pp 34–41. 10.1016/j.addr.2016.04.029.

(42) Lipinski, C. A. Drug-like Properties and the Causes of Poor Solubility and Poor Permeability. J Pharmacol Toxicol Methods 2000, 44 (1), 235–249. 10.1016/S1056-8719(00)00107-6.

(43) Baell, J. B.; Holloway, G. A. New Substructure Filters for Removal of Pan Assay Interference Compounds (PAINS) from Screening Libraries and for Their Exclusion in Bioassays. J Med Chem 2010, 53 (7), 2719–2740. 10.1021/jm901137j.

(44) Capuzzi, S. J.; Muratov, E. N.; Tropsha, A. Phantom PAINS : Problems with the Utility of Alerts for Pan - Assay INterference CompoundS. 2017. 10.1021/acs.jcim.6b00465.

(45) Milletti, F.; Storchi, L.; Sforna, G.; Cruciani, G. New and Original PKa Prediction Method Using Grid Molecular Interaction Fields. J Chem Inf Model 2007, 47 (6), 2172–2181. 10.1021/ci700018y.

(46) OMEGA v.4.1.2.0. Santa Fe, NM, USA. OpenEye Scientific Software. OpenEye Scientific Software: Santa Fe, NM, USA.

(47) Hawkins, P. C. D.; Nicholls, A. Conformer Generation with OMEGA: Learning from the Data Set and the Analysis of Failures. J Chem Inf Model 2012, 52 (11), 2919–2936. 10.1021/ci300314k.

(48) Hawkins, P. C. D.; Skillman, A. G.; Warren, G. L.; Ellingson, B. A.; Stahl, M. T. Conformer Generation with OMEGA: Algorithm and Validation Using High Quality Structures from the Protein Databank and Cambridge Structural Database - Journal of Chemical Information and Modeling (ACS Publications). 2010, 11, 572–584. 10.1021/ci100031x.

(49) ROCS v.3.5.0.2. Santa Fe, NM, USA. OpenEye Scientific Software. OpenEye Scientific Software: Santa Fe, NM, USA.

(50) Hawkins, P. C. D.; Skillman, A. G.; Nicholls, A. Comparison of Shape-Matching and Docking as Virtual Screening Tools. J Med Chem 2007, 50 (1), 74–82. 10.1021/jm0603365.

(51) O’Boyle, N. M.; Banck, M.; James, C. A.; Morley, C.; Vandermeersch, T.; Hutchison, G. R. Open Babel: An Open Chemical Toolbox. J Cheminform 2011, 3 (10), 33. 10.1186/1758-2946-3-33.

(52) QUACPAC v.2.1.3.0. Santa Fe, NM, USA. OpenEye Scientific Software.

(53) Wolber, G.; Langer, T. LigandScout: 3-D Pharmacophores Derived from Protein-Bound Ligands and Their Use as Virtual Screening Filters. J Chem Inf Model 2005, 45 (1), 160–169. 10.1021/ci049885e.

(54) Maggiora, G.; Vogt, M.; Stumpfe, D.; Bajorath, J. Molecular Similarity in Medicinal Chemistry. Journal of Medicinal Chemistry. American Chemical Society April 24, 2014, pp 3186–3204. 10.1021/jm401411z.

(55) Rogers, D. J.; Tanimoto, T. T. A Computer Program for Classifying Plants. Science (1979) 1960, 132, 1115–1118.

(56) R Core Team. R Foundation for Statistical Computing. R: A language and environment for statistical computing.: Vienna, Austria 2020.

(57) Verdonk, M. L.; Cole, J. C.; Hartshorn, M. J.; Murray, C. W.; Taylor, R. D. Improved Protein – Ligand Docking Using GOLD. Proteins Structure Function And Bioinformatics 2003, 623 (January), 609–623. 10.1002/prot.10465.

(58) Korb, O.; Stützle, T.; Exner, T. E. Empirical Scoring Functions for Advanced Protein-Ligand Docking with PLANTS. J Chem Inf Model 2009, 49 (1), 84–96. 10.1021/ci800298z.

(59) The PyMOL Molecular Graphics System v.2.5.0. Schrödinger LCC. Schrödinger LLC.

(60) Gasteiger E., H. C., G. A., D. S., W. M. R., A. R. D., B. A. Protein Identification and Analysis Tools on the Expasy Server. In The Proteomics Protocols Handbook; Walker, J. M., Ed.; Humana Totowa, NJ, 2005; pp 571–607.

(61) Henrich, S.; Salo-Ahen, O. M. H.; Huang, B.; Rippmann, F.; Cruciani, G.; Wade, R. C. Computational Approaches to Identifying and Characterizing Protein Binding Sites for Ligand Design. Journal of Molecular Recognition 2010, 23 (2), 209–219. 10.1002/jmr.984.

(62) Kokh, D. B.; Richter, S.; Henrich, S.; Czodrowski, P.; Rippmann, F.; Wade, R. C. TRAPP: A Tool for Analysis of Transient Binding Pockets in Proteins. J Chem Inf Model 2013, 53, 1235–1252. 10.1021/ci4000294.

(63) Proschak, E.; Heitel, P.; Kalinowsky, L.; Merk, D. Opportunities and Challenges for Fatty Acid Mimetics in Drug Discovery. Journal of Medicinal Chemistry. American Chemical Society July 13, 2017, pp 5235–5266. 10.1021/acs.jmedchem.6b01287.

(64) Leão, R. P.; Cruz, J. V.; da Costa, G. V.; Cruz, J. N.; Ferreira, E. F. B.; Silva, R. C.; de Lima, L. R.; Borges, R. S.; Dos Santos, G. B.; Santos, C. B. R. Identification of New Rofecoxib-Based Cyclooxygenase-2 Inhibitors: A Bioinformatics Approach. Pharmaceuticals 2020, 13 (9), 1–26. 10.3390/ph13090209.

(65) Markt, P.; Petersen, R. K.; Flindt, E. N.; Kristiansen, K.; Kirchmair, J.; Spitzer, G.; Distinto, S.; Schuster, D.; Wolber, G.; Laggner, C.; Langer, T. Discovery of Novel PPAR Ligands by a Virtual Screening Approach Based on Pharmacophore Modeling, 3D Shape, and Electrostatic Similarity Screening. J Med Chem 2008, 51 (20), 6303–6317. 10.1021/jm800128k.

(66) Lu, I. L.; Huang, C. F.; Peng, Y. H.; Lin, Y. T.; Hsieh, H. P.; Chen, C. T.; Lien, T. W.; Lee, H. J.; Mahindroo, N.; Prakash, E.; Yueh, A.; Chen, H. Y.; Goparaju, C. M. V.; Chen, X.; Liao, C. C.; Chao, Y. S.; Hsu, J. T. A.; Wu, S. Y. Structure-Based Drug Design of a Novel Family of PPARγ Partial Agonists: Virtual Screening, X-Ray Crystallography, and in Vitro/in Vivo Biological Activities. J Med Chem 2006, 49 (9), 2703–2712. 10.1021/jm051129s.

(67) Kirchmair, J.; Distinto, S.; Markt, P.; Schuster, D.; Spitzer, G. M.; Liedl, K. R.; Wolber, G. How to Optimize Shape-Based Virtual Screening: Choosing the Right Query and Including Chemical Information. J Chem Inf Model 2009, 49 (3), 678–692. 10.1021/ci8004226.

(68) Rörsch, F.; Wobst, I.; Zettl, H.; Schubert-Zsilavecz, M.; Grösch, S.; Geisslinger, G.; Schneider, G.; Proschak, E. Nonacidic Inhibitors of Human Microsomal Prostaglandin Synthase 1 (MPGES 1) Identified by a Multistep Virtual Screening Protocol. J Med Chem 2010, 53 (2), 911–915. 10.1021/jm9012505.

(69) Schuster, D.; Markt, P.; Grienke, U.; Mihaly-Bison, J.; Binder, M.; Noha, S. M.; Rollinger, J. M.; Stuppner, H.; Bochkov, V. N.; Wolber, G. Pharmacophore-Based Discovery of FXR Agonists. Part I: Model Development and Experimental Validation. Bioorg Med Chem 2011, 19 (23), 7168–7180. 10.1016/j.bmc.2011.09.056.

(70) Temml, V.; Voss, C. V.; Dirsch, V. M.; Schuster, D. Discovery of New Liver X Receptor Agonists by Pharmacophore Modeling and Shape-Based Virtual Screening. J Chem Inf Model 2014, 54 (2), 367–371. 10.1021/ci400682b.

(71) Hofmann, B.; Franke, L.; Proschak, E.; Tanrikulu, Y.; Schneider, P.; Steinhilber, D.; Schneider, G. Scaffold-Hopping Cascade Yields Potent Inhibitors of 5-Lipoxygenase. ChemMedChem 2008, 3 (10), 1535–1538. 10.1002/cmdc.200800153.

(72) Viviani, L. G.; Piccirillo, E.; Ulrich, H.; Amaral, A. T. D. Virtual Screening Approach for the Identification of Hydroxamic Acids as Novel Human Ecto-5′-Nucleotidase Inhibitors. J Chem Inf Model 2020, 60 (2), 621–630. 10.1021/acs.jcim.9b00884.

(73) Mooij, W. T. M.; Verdonk, M. L. General and Targeted Statistical Potentials for Protein-Ligand Interactions. *Proteins: Structure*, Function and Genetics 2005, 61 (2), 272–287. 10.1002/prot.20588.

(74) Jones, G.; Willett, P.; Glen, R. C.; Leach, A. R.; Taylor, R. Development and Validation of a Genetic Algorithm for Flexible Docking. J Mol Biol 1997, 267 (3), 727–748. 10.1006/jmbi.1996.0897.

(75) Feng, B. Y.; Shoichet, B. K. A Detergent-Based Assay for the Detection of Promiscuous Inhibitors. Nat Protoc 2006, 1 (2), 550–553. 10.1038/nprot.2006.77.

(76) McGovern, S. L.; Caselli, E.; Grigorieff, N.; Shoichet, B. K. A Common Mechanism Underlying Promiscuous Inhibitors from Virtual and High-Throughput Screening. J Med Chem 2002, 45 (8), 1712–1722. 10.1021/jm010533y.

(77) McGovern, S. L.; Helfand, B. T.; Feng, B.; Shoichet, B. K. A Specific Mechanism of Nonspecific Inhibition. J Med Chem 2003, 46 (20), 4265–4272. 10.1021/jm030266r.

(78) Aldrich, C.; Bertozzi, C.; Georg, G. I.; Kiessling, L.; Lindsley, C.; Liotta, D.; Merz, K. M.; Schepartz, A.; Wang, S. The Ecstasy and Agony of Assay Interference Compounds. Journal of Medicinal Chemistry. American Chemical Society March 23, 2017, pp 2165–2168. 10.1021/acs.jmedchem.7b00229.

(79) Viviani, L. G.; Piccirillo, E.; Cheffer, A.; de Rezende, L.; Maria Carmona-Ribeiro, A.; T-do Amaral, A. Identifying Aggregators on a Virtual Screening Search for Potential Human Ecto-5’-Nucleotidase Inhibitors 4; 2018; Vol. 23. www.mdpi.com/journal/molecules.

(80) Lipinski, C. A.; Lombardo, F.; Dominy, B. W.; Feeney, P. J. Experimental and Computational Approaches to Estimate Solubility and Permeability in Drug Discovery and Develop Ment Settings. Adv Drug Deliv Rev 1997, 23, 3–25. 10.1016/S0169-409X(00)00129-0.

(81) Tetko, I. V.; Tanchuk, V. Y.; Kasheva, T. N.; Villa, A. E. P. Estimation of Aqueous Solubility of Chemical Compounds Using E-State Indices. J Chem Inf Comput Sci 2001, 41 (6), 1488–1493. 10.1021/ci000392t.

(82) Llompart, P.; Minoletti, C.; Baybekov, S.; Horvath, D.; Marcou, G.; Varnek, A. Will We Ever Be Able to Accurately Predict Solubility? Sci Data 2024, 11 (1). 10.1038/s41597-024-03105-6.

(83) Daina, A.; Michielin, O.; Zoete, V. SwissADME: A Free Web Tool to Evaluate Pharmacokinetics, Drug-Likeness and Medicinal Chemistry Friendliness of Small Molecules. Sci Rep 2017, 7. 10.1038/srep42717.

(84) Pesaresi, A. Mixed and Non-Competitive Enzyme Inhibition: Underlying Mechanisms and Mechanistic Irrelevance of the Formal Two-Site Model. J Enzyme Inhib Med Chem 2023, 38 (1). 10.1080/14756366.2023.2245168.

(85) Blat, Y. Non-Competitive Inhibition by Active Site Binders. Chemical Biology and Drug Design. June 2010, pp 535–540. 10.1111/j.1747-0285.2010.00972.x.

(86) Buker, S. M.; Boriack-Sjodin, P. A.; Copeland, R. A. Enzyme–Inhibitor Interactions and a Simple, Rapid Method for Determining Inhibition Modality. SLAS Discovery 2019, 24 (5), 515–522. 10.1177/2472555219829898.

(87) Scior, T.; Bender, A.; Tresadern, G.; Medina-Franco, J. L.; Martínez-Mayorga, K.; Langer, T.; Cuanalo-Contreras, K.; Agrafiotis, D. K. Recognizing Pitfalls in Virtual Screening: A Critical Review. J Chem Inf Model 2012, 52 (4), 867–881. 10.1021/ci200528d.

(88) Koutsoukas, A.; Paricharak, S.; Galloway, W.; Spring, D. R.; Ijzerman, A. P.; Glen, R.; Marcus, D.; Bender, A. How Diverse Are Diversity Assessment MethodsA Comparative Analysis and Benchmarking of Molecular Descriptor Space Unilever Centre for Molecular Science Informatics, Department of Chemistry, University of Division of Medicinal Chemistry, Leiden Acade. 2013.

(89) Nicholls, A.; McGaughey, G. B.; Sheridan, R. P.; Good, A. C.; Warren, G.; Mathieu, M.; Muchmore, S. W.; Brown, S. P.; Grant, J. A.; Haigh, J. A.; Nevins, N.; Jain, A. N.; Kelley, B. Molecular Shape and Medicinal Chemistry: A Perspective. Journal of Medicinal Chemistry. May 27, 2010, pp 3862–3886. 10.1021/jm900818s.

(90) Daina, A.; Michielin, O.; Zoete, V. SwissADME: A Free Web Tool to Evaluate Pharmacokinetics, Drug-Likeness and Medicinal Chemistry Friendliness of Small Molecules. Sci Rep 2017, 7. 10.1038/srep42717.

(91) Heiden, W.; Moeckel, G.; Brickmann, J. A New Approach to Analysis and Display of Local Lipophilicity/ Hydrophilicity Mapped on Molecular Surfaces; 1993; Vol. 7.

(92) Connolly, M. Solvent-Accessible Surfaces of Proteins and Nucleic Acids. Science (1979) 1983, 221 (4612), 709–713. 10.1126/science.6879170.

