## Supporting Information for "Identification of novel human 15-lipoxygenase-2 (h15-LOX-2) inhibitors using a virtual screening approach"

**Table S1.** RMSD values calculated between the pose of the substrate mimic inhibitor C8E4 observed in the crystal structure (PDB code: 4NRE)[1] and the two best-scored docking poses (poses 1 and 2).

| **Pose (ranked)** | **RMSD values(*) (Å)** | | | |
|---|---|---|---|---|
| | **ASP** | **ChemPLP** | **ChemScore** | **GoldScore** |
| 1 | 2.65 | 2.75 | 7.06 | 2.07 |
| 2 | 4.02 | 2.82 | 6.93 | 3.89 |
| Average (1 and 2) | 3.34 | 2.79 | 7.00 | 2.98 |

(*) Calculated between the docking pose and the pose observed in the crystal structure (PDB code: 4NRE).[1]

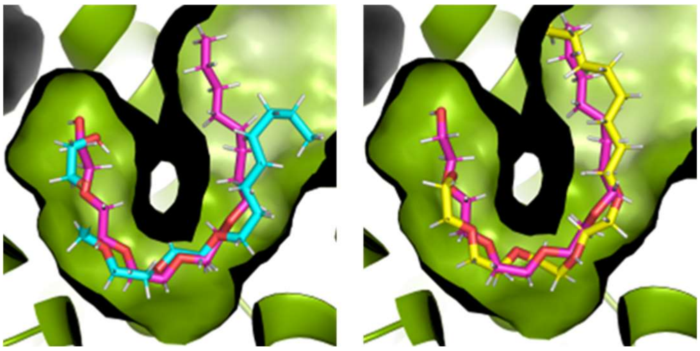

**Figure S1.** Superposition between the crystallographic pose observed for the inhibitor C8E4 in the 15-LOX-2-C8E4 complex (PDB code: 4NRE[1], magenta sticks) and the respective best-scored docking solutions (pose 1) obtained using ChemPLP (cyan sticks, left) and GoldScore (yellow sticks, right) scoring functions. The figure was prepared using PyMOL.

---

[1] Kobe et al. *J. Biol. Chem.*, 289, 8562-8569, 2014.

**Table S2.** Structures and physicochemical properties of the compounds selected as potential h15-LOX-2 inhibitors from the ZINC-Curated database by the proposed VS protocol.

| Name (Ref. and/or ZINC ID) | Structure | Molecular formula | Molecular weight (g.mol$^{-1}$) | LogP[(*)] |
|---|---|---|---|---|
| **01** ZINC63152202 | 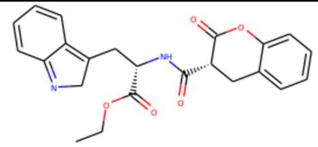 | $C_{23}H_{22}N_2O_5$ | 406.4 | 0.69 |
| **02** ZINC64045009 | 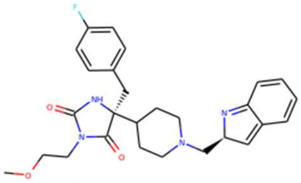 | $C_{27}H_{31}FN_4O_3$ | 479.6 | 1.5 |
| **03** ZINC14989654 | 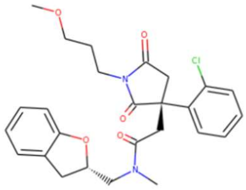 | $C_{26}H_{29}ClN_2O_5$ | 485.0 | 3.2 |
| **04** ZINC21018084 | 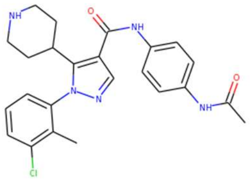 | $C_{24}H_{26}ClN_5O_2$ | 453.0 | 4.5 |
| **05** ZINC63667415 | 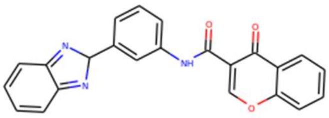 | $C_{23}H_{15}N_3O_3$ | 381.4 | 3.0 |
| **06** ZINC09445447 | 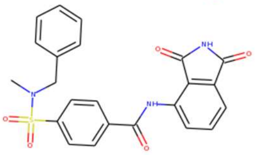 | $C_{23}H_{19}N_3O_5S$ | 449.5 | 2.6 |
| **07** ZINC10187184 | 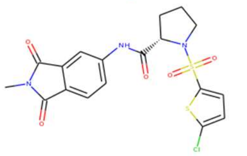 | $C_{18}H_{16}ClN_3O_5S_2$ | 453.9 | 2.4 |
| **08** ZINC17148759 | 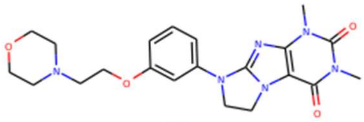 | $C_{21}H_{26}N_6O_4$ | 426.5 | 0.29 |
| **09** ZINC23138421 | 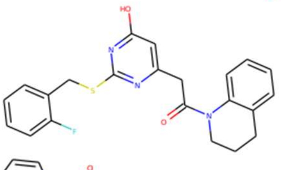 | $C_{22}H_{20}FN_3O_2S$ | 409.5 | 4.1 |
| **10** ZINC00794703 | 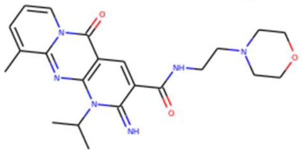 | $C_{22}H_{28}N_6O_3$ | 425.5 | 1.1 |
| **11** ZINC02395677 | 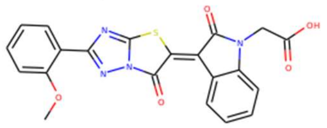 | $C_{21}H_{14}N_4O_5S$ | 433.4 | 1.2 |

| | | | | |
|---|---|---|---|---|
| **12** ZINC63362107 | 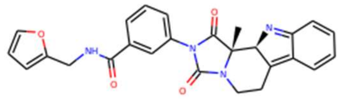 | $C_{26}H_{22}N_4O_4$ | 454.5 | 2.0 |
| **13** ZINC18202958 | 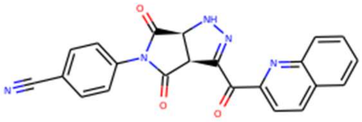 | $C_{22}H_{13}N_5O_3$ | 395.4 | 1.8 |
| **14** ZINC32124366 | 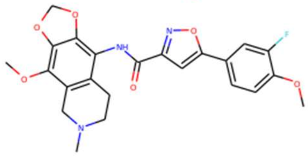 | $C_{23}H_{22}FN_3O_6$ | 456.5 | 3.5 |
| ZINC79045996 | 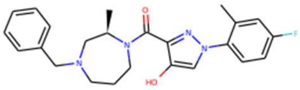 | $C_{24}H_{27}FN_4O_2$ | 423.5 | 3.8 |
| ZINC77939079 | 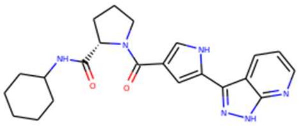 | $C_{22}H_{26}N_6O_2$ | 406.5 | 3.0 |
| ZINC14012965 | 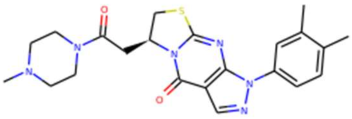 | $C_{22}H_{26}N_6O_2S$ | 439.6 | 2.0 |
| ZINC39733003 | 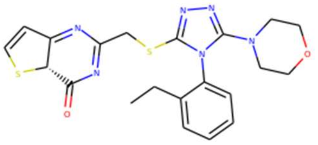 | $C_{21}H_{22}N_6O_2S_2$ | 454.6 | 2.8 |
| ZINC64622398 | 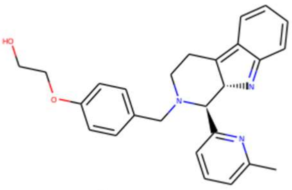 | $C_{26}H_{27}N_3O_2$ | 414.5 | 2.6 |
| ZINC13109340 | 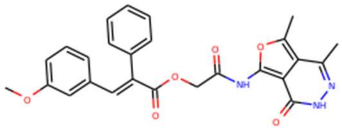 | $C_{26}H_{23}N_3O_6$ | 473.5 | 3.9 |
| ZINC33010296 | 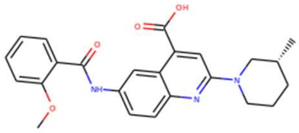 | $C_{24}H_{25}N_3O_4$ | 418.5 | 4.4 |
| ZINC33260851 | 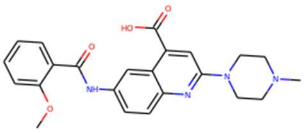 | $C_{23}H_{24}N_4O_4$ | 420.5 | 2.9 |
| ZINC15269234 | 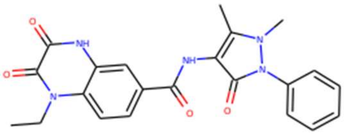 | $C_{22}H_{21}N_5O_4$ | 419.4 | 1.8 |
| ZINC72465538 | 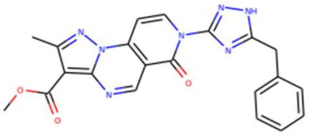 | $C_{21}H_{17}N_7O_3$ | 414.4 | 1.8 |

| | | | | |
|---|---|---|---|---|
| ZINC19418460 | 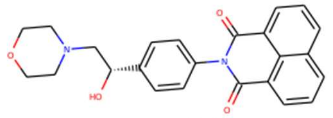 | $C_{24}H_{22}N_2O_4$ | 402.5 | 3.0 |
| ZINC09120398 | 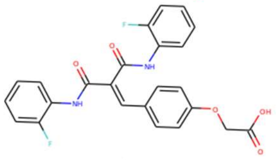 | $C_{24}H_{18}F_2N_2O_5$ | 451.4 | 4.1 |
| ZINC39467137 | 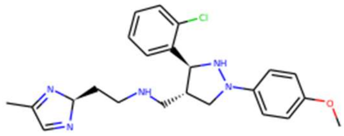 | $C_{23}H_{28}ClN_5O$ | 427.0 | 3.9 |
| ZINC02323981 | 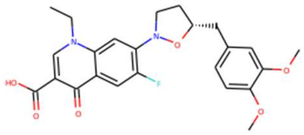 | $C_{24}H_{25}FN_2O_6$ | 455.5 | 3.6 |
| ZINC11785568 | 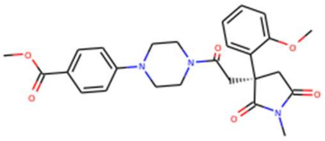 | $C_{26}H_{29}N_3O_6$ | 479.5 | 1.8 |
| ZINC67554839 | 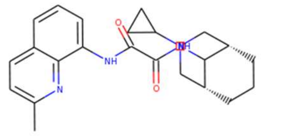 | $C_{23}H_{28}N_4O_2$ | 393.5 | 2.9 |
| ZINC20792548 | 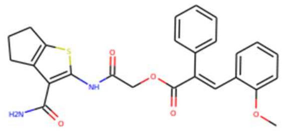 | $C_{26}H_{24}N_2O_5S$ | 476.6 | 4.1 |
| ZINC63667383 | 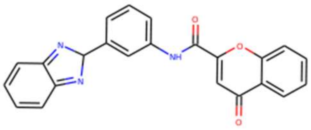 | $C_{23}H_{15}N_3O_3$ | 381.4 | 3.0 |
| ZINC64007007 | 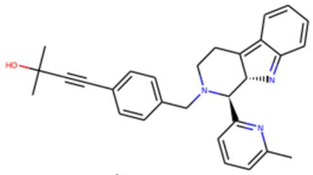 | $C_{29}H_{29}N_3O$ | 436.6 | 3.3 |
| ZINC12035964 | 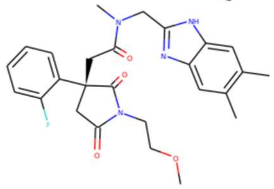 | $C_{26}H_{29}FN_4O_4$ | 480.5 | 3.0 |
| ZINC11783806 | 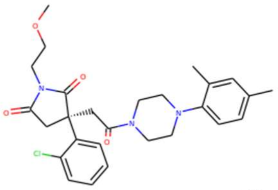 | $C_{27}H_{32}ClN_3O_4$ | 498.0 | 3.3 |
| ZINC64031717 | 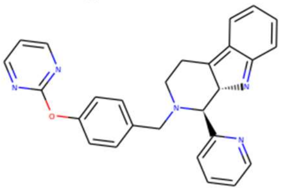 | $C_{27}H_{23}N_5O$ | 434.5 | 3.5 |

| | | | | |
|---|---|---|---|---|
| ZINC69046714 | 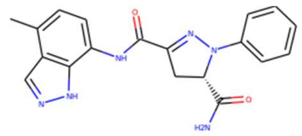 | $C_{19}H_{18}N_6O_2$ | 362.4 | 1.9 |
| ZINC38631038 | 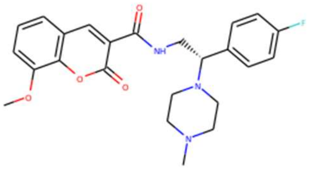 | $C_{24}H_{26}FN_3O_4$ | 440.5 | 2.7 |
| ZINC12037832 | 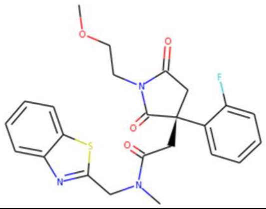 | $C_{24}H_{24}FN_3O_4S$ | 469.5 | 3.1 |

(*) LogP values were taken from the ZINC database.

Compound 01

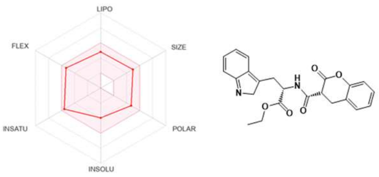


Compound 02

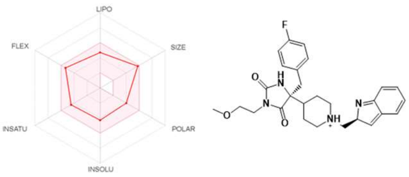


Compound 03

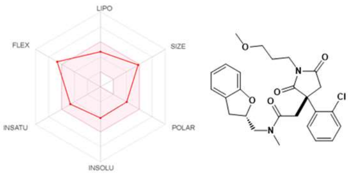


Compound 04

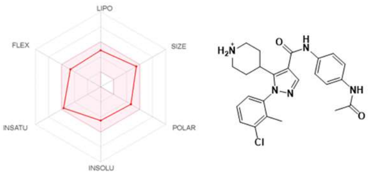


Compound 05

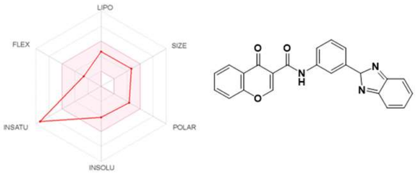


Compound 06

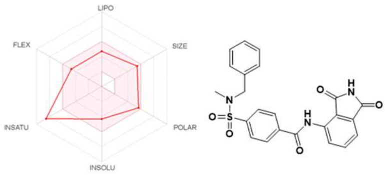


Compound 07

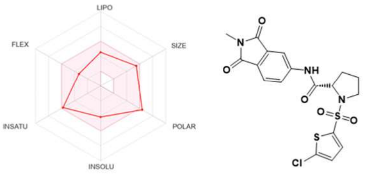


Compound 08

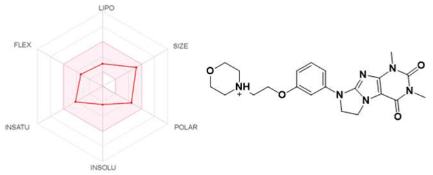


Compound 09

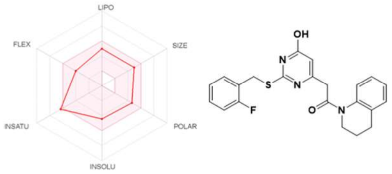


Compound 10

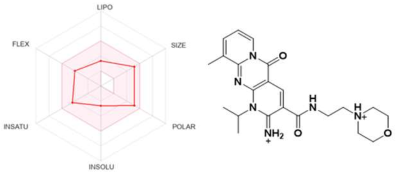


Compound 11

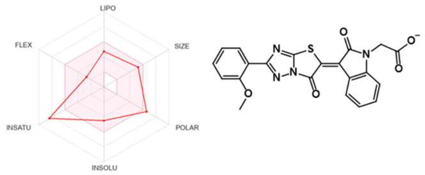


Compound 12

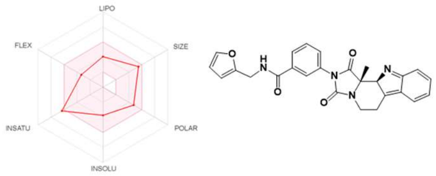


Compound 13

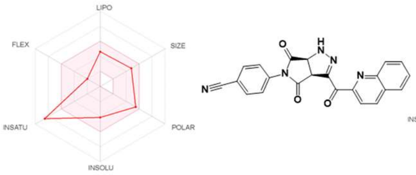


Compound 14

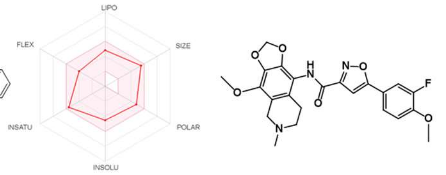


**Figure S2. SwissADME[1] "bioavailability radar" plots for the compounds selected as h15-LOX-2 inhibitor candidates from the ZINC-Curated database and that have been acquired for enzymatic inhibitory assays.** The colored zone in the diagrams represents the suitable physicochemical space for oral bioavailability, according to the following parameter limits: LIPO (lipophilicity): XLOGP3 between −0.7 and +5.0; SIZE: molecular weight between 150 and 500 g.mol$^{-1}$; POLAR (polarity): TPSA between 20 and 130 Å$^2$; INSOLU (solubility): LogS not higher than 6; INSATU (degree of saturation): fraction of carbons with sp$^3$ hybridization between 0.25 and 1.0; and FLEX (flexibility): no more than 9 rotatable bonds.

---

[1] Daina, A., Michielin, O., and Zoete, V. (2017). SwissADME: A free web tool to evaluate pharmacokinetics, drug-likeness and medicinal chemistry friendliness of small molecules. *Scientific Reports* 7. DOI: 10.1038/srep42717.

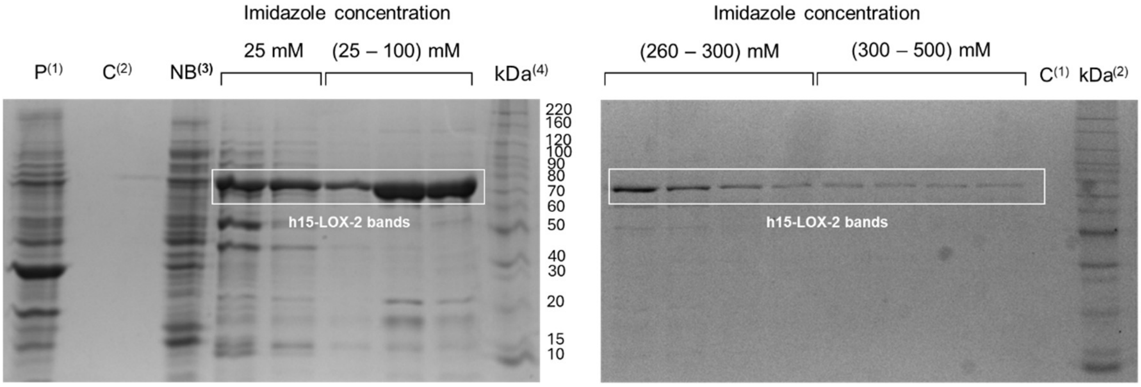


**Figure S3. Representative results of the expression and purification of h15-LOX-2 (SDS-PAGE analysis of the collected peaks).** (1) P: Cell pellet; (2) C: Control (loading buffer); (3) NB: Lysate supernatant not-bound to the $Ni^{2+}$-NTA affinity column; (4) Molecular weight markers: BenchMark™ Protein Ladder (*Invitrogen*).

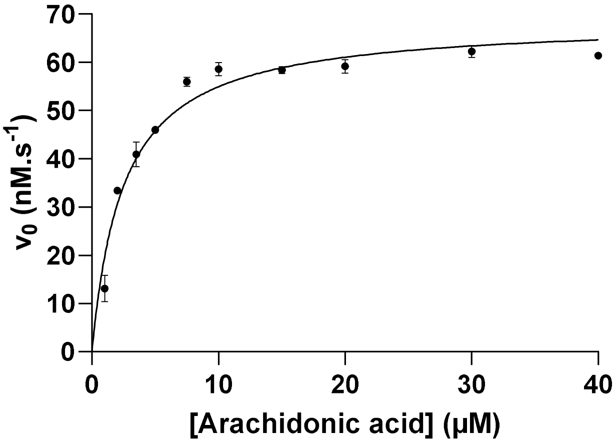


**Figure S4.** Plot of initial velocities of h15-LOX-2-catalyzed reaction as a function of arachidonic acid concentrations. A Michaelis-Menten model was used to fit the experimental data. Assays were carried out in a total volume of 300 µL, containing Tris-HCl (25 mM, pH = 8.0), NaCl (250 mM), h15-LOX-2 (120 nM), Triton X-100 (0.017% v/v), and arachidonic acid (1.0 - 40 µM). Data represent the average ± S.E.M. of experiments performed twice and at least in duplicates. The figure was prepared using GraphPad Prism.

**Table S3.** Kinetic parameters calculated for h15-LOX-2-catalyzed reaction, using arachidonic acid as a substrate.[1]

| Kinetic parameter | Experimentally obtained value | Reference value (literature)[2] |
|---|---|---|
| $K_M$ [95% C.I.][3] or ± S.D.[4] (μM) | 2.51 [2.06; 3.03] | 1.9 ± 0.37 |
| $V_{max}$ [95% C.I.][3] ($nM{\cdot}s^{-1}$) | 68.70 [65.35; 72.26] | - |
| $k_{cat}$ ($s^{-1}$) ± S.D.[4] | 0.57 | 0.6 ± 0.02 |

(1) Data represent the averages from two independent experiments performed in duplicates. Assays were carried out in a total volume of 300 μL, containing Tris-HCl (25 mM, pH = 8.0), NaCl (250 mM), h15-LOX-2 (120 nM), Triton X-100 (0.017% v/v), and arachidonic acid (1.0 - 40 μM).

(2) Reference: Kobe et al., *J. Biol. Chem.*, 289, 12, 2014.

(3) C.I.: confidence interval.

(4) S.D.: standard deviation.

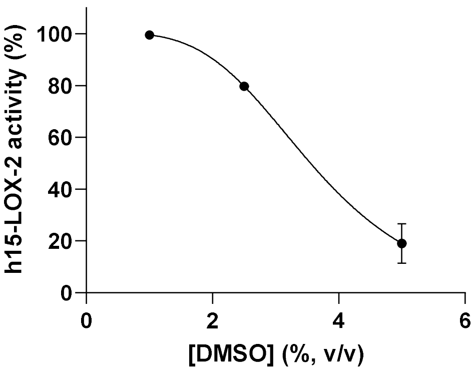


**Figure S5.** Effect of DMSO on the h15-LOX-2 activity. Assays were carried out in a total volume of 300 μL, containing Tris-HCl (25 mM, pH = 8.0), NaCl (250 mM), h15-LOX-2 (120 nM), Triton X-100 (0.01% v/v), arachidonic acid (25 μM), and DMSO (0 - 5.0%, v/v). Data represent the average ± S.E.M. of experiments performed in duplicates. The figure was prepared using GraphPad Prism.

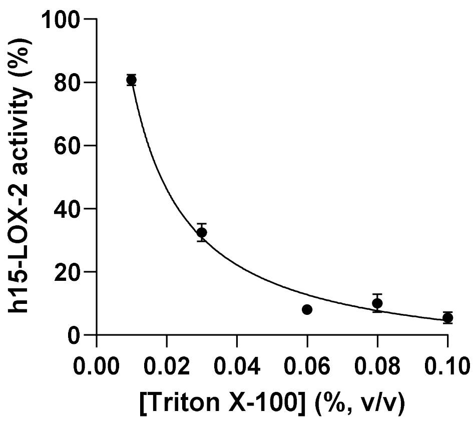


**Figure S6.** Effect of Triton X-100 on the h15-LOX-2 activity. Assays were carried out in a total volume of 300 µL, containing Tris-HCl (25 mM, pH = 8.0), NaCl (250 mM), h15-LOX-2 (420 nM), arachidonic acid (25 µM), and Triton X-100 (0 – 0.1% v/v). Data represent the average ± S.E.M. of experiments performed in duplicates. The figure was prepared using GraphPad Prism.
